# Activation of neurogenesis improves amyloid-β pathology and cognitive function through AMP kinase signaling in Alzheimer’s disease model mice

**DOI:** 10.64898/2025.12.24.696442

**Authors:** Masahiro Fukui, Takashi Kaise, Taimu Masaki, Tyler Sakamoto, Ryoichiro Kageyama

**Author notes:** Correspondence: Ryoichiro Kageyama, RIKEN Center for Biosystems Dynamics Research, Kobe 650-0047, Japan.

## Abstract

Adult hippocampal neurogenesis declines with aging and in neurological disorders, leading to cognitive impairment. We previously demonstrated that a treatment of inducing *Plagl2*, a zinc finger transcription factor gene, and antagonizing *Dyrk1a*, a gene associated with Down syndrome, referred to as iPaD, can functionally rejuvenate aged neural stem cells (NSCs), thereby enhancing neurogenesis and improving cognition in aged mice. Here, we found that NSC-specific iPaD treatment effectively activated neurogenesis, reduced amyloid-β deposition, and improved cognitive function in Alzheimer’s disease model mice. Transcriptomic analysis revealed widespread significant changes in gene expression in the hippocampus following iPaD treatment. The upregulated genes included those associated with the activation of astrocytes and microglia involved in amyloid-β clearance, while the downregulated genes included several that are upregulated in Alzheimer’s disease patients, but whose roles in disease progression remain unclear. Among the latter genes, knockdown of *Prkag2*, a gene encoding protein kinase AMP-activated non-catalytic subunit gamma 2, in the hippocampus most effectively enhanced neurogenesis and reduced amyloid-β accumulation. Notably, both iPaD treatment and *Prkag2* knockdown activated AMP-activated protein kinase signaling, thereby upregulating genes involved in autophagy and cellular homeostasis. These results suggest that *Prkag2* may represent a promising therapeutic target for neurodegenerative diseases, including Alzheimer’s disease.

## Introduction

Neural stem cells (NSCs) are present in the adult hippocampal dentate gyrus of various species including mice and humans (Gonçalves et al. 2016; Moreno-Jiménez et al. 2019; Tobin et al. 2019; Zhou et al. 2022). Adult NSCs are mostly quiescent/dormant but are occasionally activated to start proliferating and generate new neurons, a phenomenon called adult neurogenesis (Gonçalves et al. 2016). Adult-born hippocampal neurons play an important role in learning, memory, and cognition, but adult neurogenesis declines with aging (Imayoshi et al. 2008; Lugert et al. 2010; Encinas et al. 2011). The activation of neurogenesis can ameliorate age-related cognitive dysfunction and neurodegenerative disorders (Benraiss et al. 2013; Choi et al. 2018; Díaz-Moreno et al. 2018), but it usually leads to the exhaustion of NSCs, thereby terminating neurogenesis prematurely (Encinas et al. 2011; Sueda et al. 2019; Zhang et al. 2019). Similarly, adult neurogenesis is impaired in Alzheimer’s disease patients, the most common age-related dementia, which is characterized by amyloid-β (Aβ) deposition and subsequent neurofibrillary tangle formation, leading to neurodegeneration and cognitive impairment (Moreno-Jiménez et al. 2019; Tobin et al. 2019; Zhou et al. 2022). Aβ deposition and cognitive impairment also occur in Alzheimer’s disease model mice (Oakley et al. 2006). While enhanced neurogenesis has been shown to improve cognitive functions in Alzheimer’s disease model mice, it does not improve Aβ pathology (Choi et al. 2018; Li et al. 2023). These results suggest that the enhanced neurogenesis induced by current methods is not sufficient to treat Alzheimer’s disease, and further improved methods are required (Salta et al. 2023).

We previously demonstrated that aged NSCs that have lost their proliferative and neurogenic potential can be functionally rejuvenated by inducing *Plagl2*, a zinc finger transcription factor gene, and antagonizing *Dyrk1a*, a gene associated with Down syndrome (a genetic disorder known to accelerate aging). This combinatorial approach, which is referred to as iPaD (inducing *Plagl2* and anti-*Dyrk1a*), restores the ability of NSCs to proliferate and generate new neurons at levels comparable to those observed in the juvenile hippocampus, leading to improved cognitive function (Kaise et al., 2022). This treatment activates the oscillatory expression of the proneural gene *Ascl1*, leading to the activation of dormant NSCs and their subsequent differentiation into intermediate progenitor cells and, ultimately, mature neurons. Transcriptomic analysis revealed that iPaD treatment can rejuvenate 20-month-old NSCs to a state resembling that of 1-month-old NSCs (Kaise et al., 2022). These findings suggest that this approach can reverse NSC aging and restore sustained functional neurogenesis, offering a promising strategy for treating age-related neurological disorders.

Here, to assess the impact of the functional rejuvenation of NSCs on Alzheimer’s disease phenotypes, we delivered the iPaD lentivirus into 5xFAD mice, which are Alzheimer’s disease model mice harboring mutant human amyloid precursor protein and presenilin 1 genes under the control of the mouse Thy1 promoter (Oakley et al. 2006). iPaD treatment effectively enhanced adult neurogenesis, reduced Aβ deposition, and improved cognitive performance. To elucidate the underlying mechanisms, we conducted further analyses of 5xFAD mice and observed the upregulation of genes associated with astrocyte and microglial activation involved in Aβ clearance. Additionally, a broad downregulation of gene expression was observed; among these, knockdown of *Prkag2*, a gene upregulated in the brains of Alzheimer’s disease patients, most effectively promoted neurogenesis and reduced Aβ accumulation. Further investigation revealed that *Prkag2* knockdown activated AMP-activated protein kinase (AMPK) signaling, resulting in the upregulation of genes involved in autophagy and cellular homeostasis and ameliorating age-related neural pathology in Alzheimer’s disease model mice.

## Results

### iPaD treatment improves Alzheimer’s disease pathology in 5xFAD mice

To investigate the effect of iPaD treatment on Alzheimer’s disease, we used 5xFAD mice, which exhibit Aβ deposition and cognitive impairment (Oakley et al. 2006). In these mice, proliferative cells (MCM2^+^), including activated NSCs and intermediate progenitor cells, and immature neurons (DCX^+^;dendrite^+^) were significantly reduced in number compared to wild-type mice (Supplemental Fig. S1A-C), indicating that neurogenesis is inhibited. We injected either the control or iPaD lentivirus under the control of the Hes5 promoter, which is active in NSCs, into the hippocampal dentate gyrus of 5-month-old 5xFAD mice (Fig. 1A,B). Lentivirus-infected NSCs and their progeny became mCherry^+^ (Fig. 1C,D). When the iPaD lentivirus was injected, proliferative cells (MCM2^+^, Fig. 1C,E) and immature neurons (DCX^+^;dendrite^+^, Fig. 1D,F) were significantly increased in number at 4, 8, and 12 weeks of post infection (wpi) compared to control lentivirus injection. These results indicated that iPaD treatment effectively activates dormant NSCs, thereby enhancing neurogenesis in 5xFAD mice.

**Figure 1.**
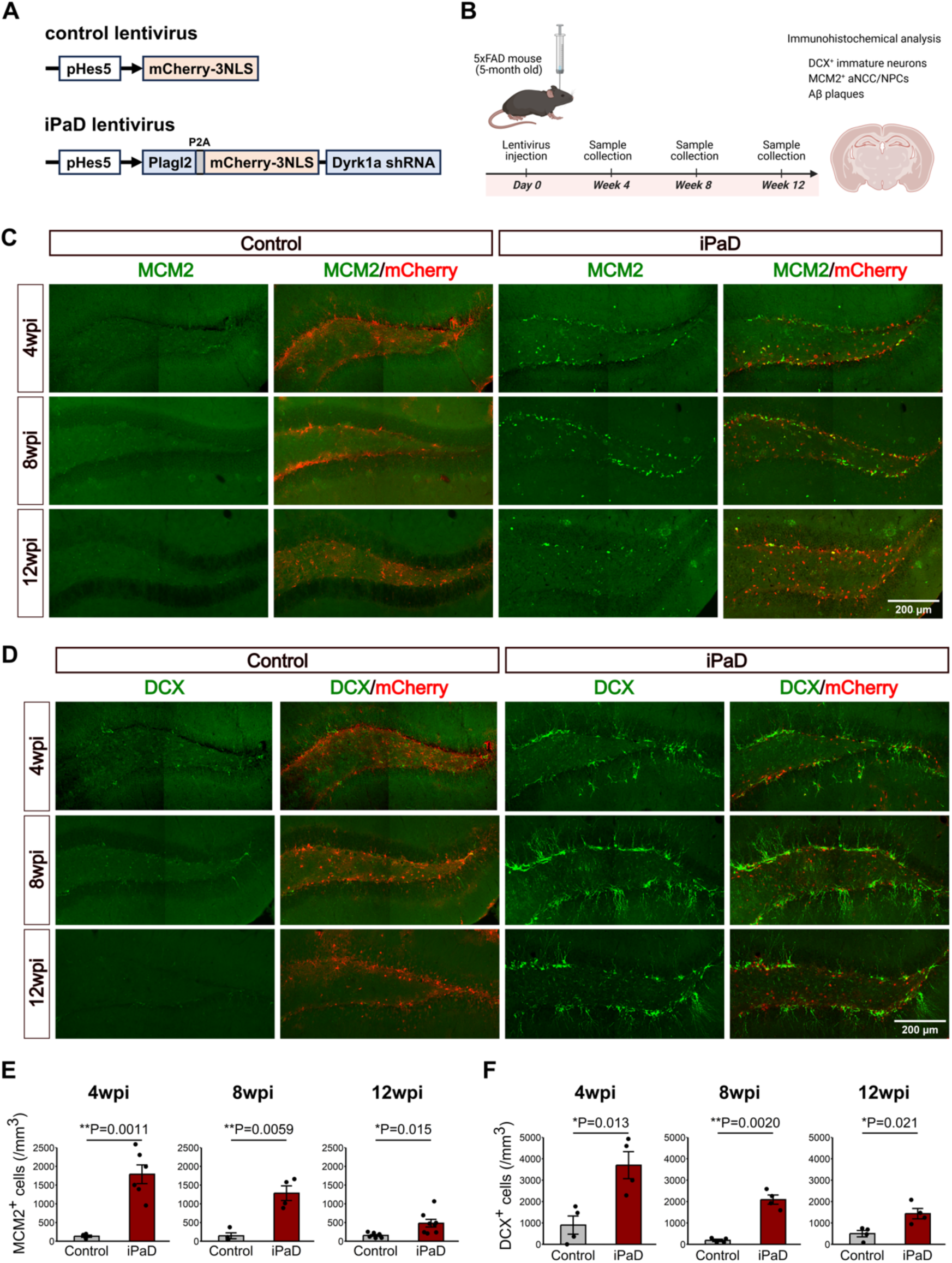
iPaD treatment enhances neurogenesis in the hippocampal dentate gyrus of 5xFAD mice. (A) Schematic structures of Hes5 promoter-driven control or iPaD lentivirus (pHes5-iPaD). (B) Experimental schedule. Virus was injected into the hippocampal dentate gyrus of 5-month-old 5xFAD mice, and brain sections were examined immunohistochemically at 4 weeks post-infection (wpi), 8 wpi, and 12 wpi. Schematic was created using BioRender. (C,D) Immunohistochemistry for MCM2^+^ activated NSCs (C) and DCX^+^ newly-born neuroblasts (D). Scale bars: 200 µm. (E,F) Quantification of MCM2^+^ activated NSCs (E) and DCX^+^ newly-born neuroblasts (F). Values are presented as the mean ± SEM. *P < 0.05, **P < 0.01, ***P < 0.001. Statistical significance was assessed using Welch’s *t* test for pairwise comparisons. (E) Control, n = 4 at 4 wpi and 8 wpi, n = 7 at 12 wpi; iPaD, n = 6 at 4 wpi, n = 4 at 8 wpi, and n = 8 at 12 wpi. (F) n = 4 per group.

In 5xFAD mice, Aβ deposition was significantly increased with aging (Supplemental Fig. S1D,E), as reported previously (Oakley et al. 2006). Notably, when the iPaD lentivirus was injected into the hippocampus of 5-month-old 5xFAD mice, Aβ deposition was significantly inhibited, compared to control (Fig. 2A-D). The inhibition of Aβ accumulation occurred not only in the granule cell layer and molecular layer, where iPaD-dependent newly born neurons reside, but also in the hilus, where no such neurons are present (Fig. 2A). These results indicated that the activation of neurogenesis by iPaD treatment improves Aβ pathology broadly in a non-cell-autonomous manner in 5xFAD mice.

**Figure 2.**
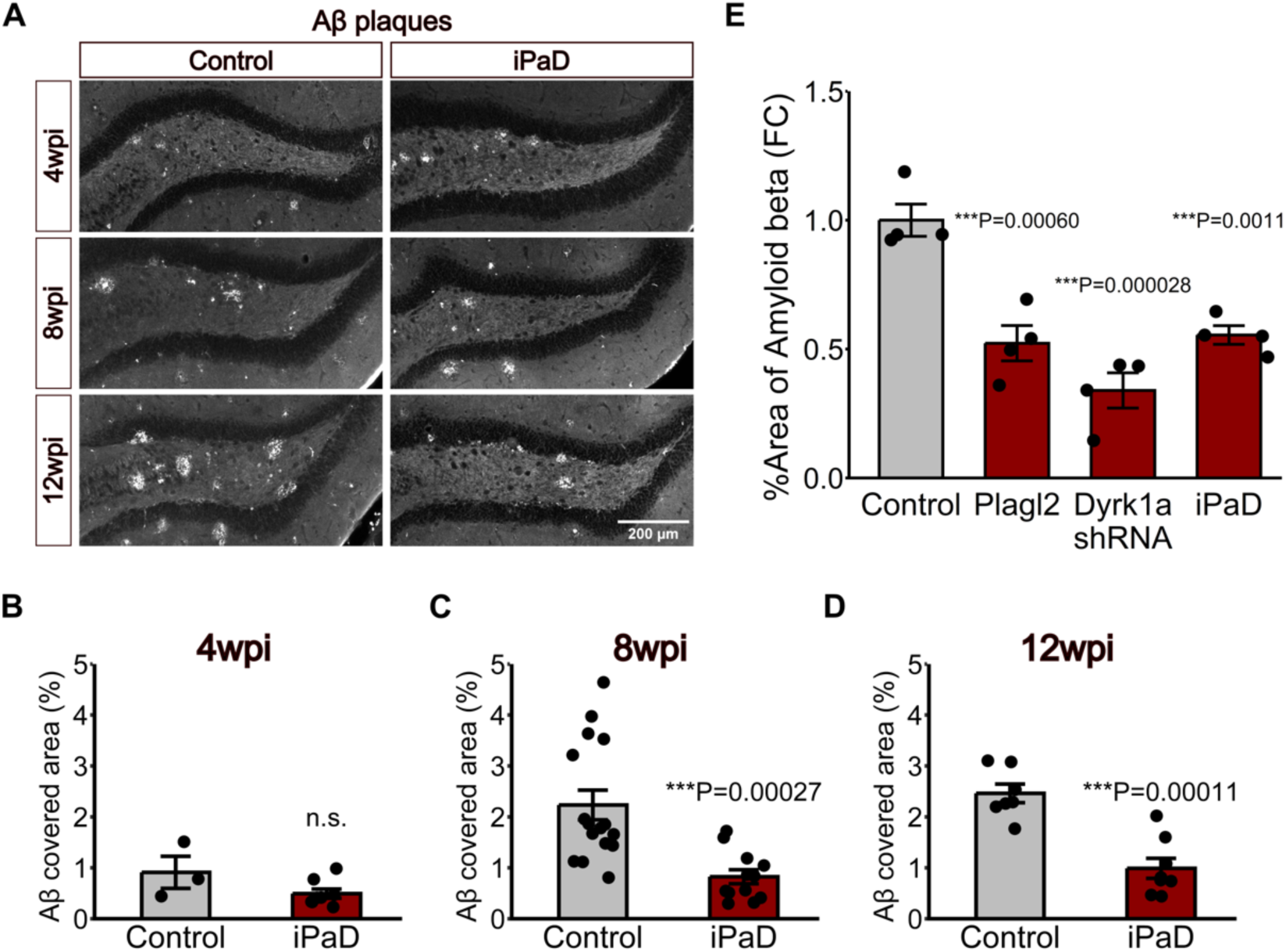
iPaD treatment decreases Aβ deposition in the hippocampal dentate gyrus of 5xFAD mice. (A-D) Hes5 promoter-driven control or iPaD lentivirus was injected into the hippocampal dentate gyrus of 5-month-old 5xFAD mice, and brain sections were examined immunohistochemically at 4 wpi, 8 wpi, and 12 wpi. Representative images (A) and quantification (B-D) of Aβ plaque area. Scale bar: 200 µm. Control, n = 4 (4 wpi), n = 16 (8 wpi), n = 7 (12 wpi); iPaD, n = 6 (4 wpi), n = 12 (8 wpi), n = 7 (12 wpi). (E) Hes5 promoter-driven control, Plagl2, Dyrk1a shRNA, or iPaD lentivirus was injected into the hippocampal dentate gyrus of 5-month-old 5xFAD mice, and Aβ plaque area was quantified at 8 wpi (n = 4 for each group). Values are presented as the mean ± SEM. *P < 0.05, **P < 0.01, ***P < 0.001. Statistical significance was assessed using Welch’s t test for pairwise comparisons, and two-way ANOVA followed by Tukey’s post hoc test for multiple group comparisons.

We previously showed that *Plagl2* overexpression alone effectively activates dormant aged NSCs, but approximately half of the progeny are neurons, while the other half are glial cells (Kaise et al. 2022). In contrast, *Dyrk1a* knockdown alone does not efficiently activate dormant NSCs, but generates mostly neurons (Kaise et al. 2022). We next examined which is involved in the reduction of Aβ deposition. Both *Plagl2* overexpression alone and *Dyrk1a* knockdown alone under the control of the Hes5 promoter effectively reduced Aβ deposition in 5xFAD mice, similar to the effect observed with iPaD treatment (Fig. 2E). These results suggest that the robust activation of neurogenesis by either method (*Plagl2* overexpression or *Dyrk1a* knockdown) may reduce Aβ deposition, thereby ameliorating Alzheimer’s disease pathology.

### iPaD treatment improves the cognitive function of 5xFAD mice

5xFAD mice reportedly exhibit memory and cognitive impairments (Oakley et al. 2006), and we confirmed such defects in these mice using the Barnes maze (Supplemental Fig. S2A-C) and fear conditioning tests (Supplemental Fig. S2D). To examine the effect of the robust activation of neurogenesis and inhibition of Aβ deposition on the cognitive function of 5xFAD mice, we injected the control or iPaD lentivirus into the hippocampus of 5-month-old 5xFAD mice and analyzed their spatial learning and memory with the Barnes maze test at 4 weeks after virus injection (Fig. 3A). Spatial learning and memory are dependent on neurogenesis in the hippocampal dentate gyrus (Imayoshi et al., 2008), and we found that iPaD lentivirus-infected 5xFAD mice reached the target hole with a shorter latency, shorter distance, and fewer errors than those infected with the control lentivirus (Fig. 3B–E). These data suggest that injection of the iPaD lentivirus improves the spatial memory of 5xFAD mice in the Barnes maze test.

**Figure 3.**
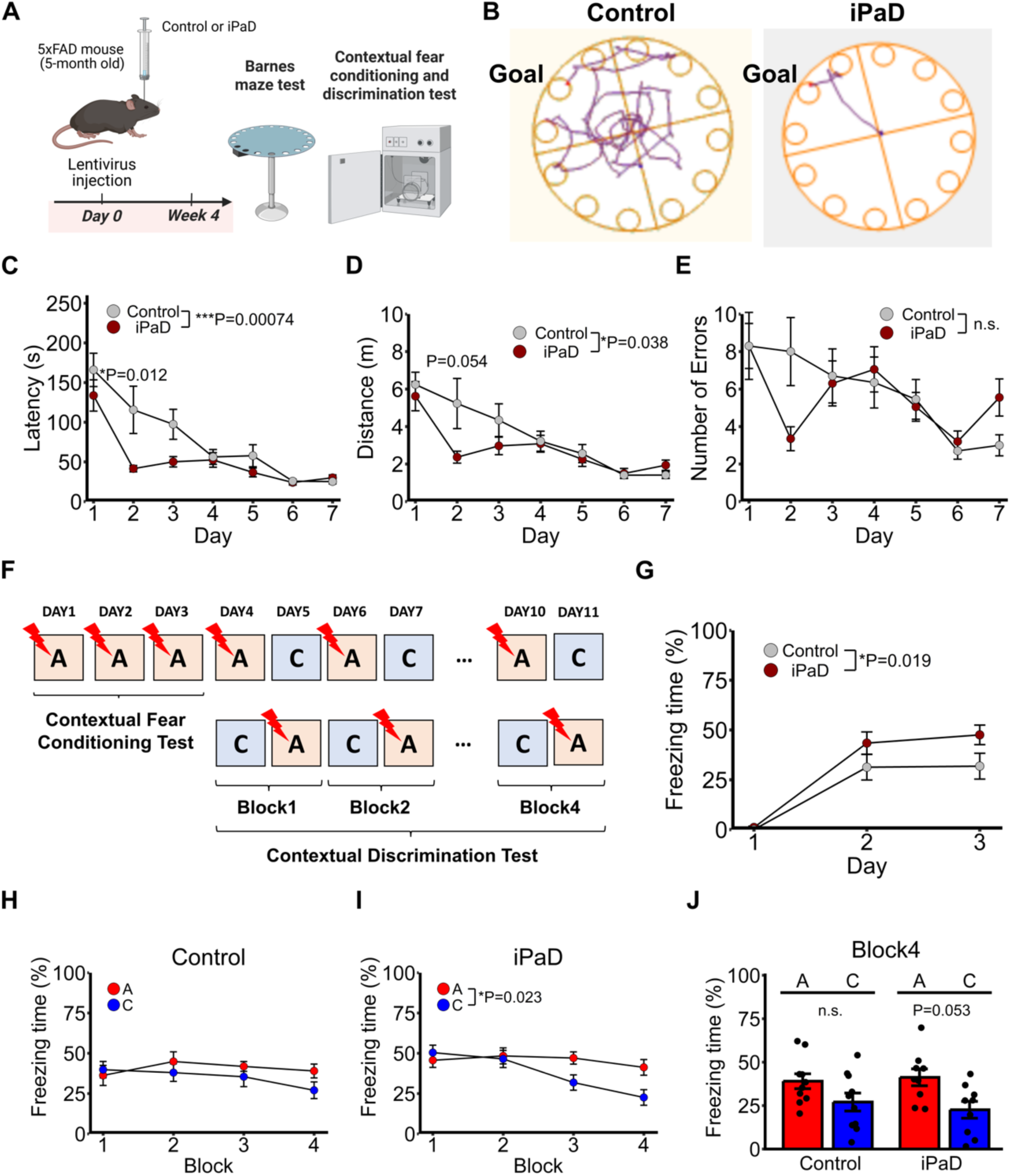
iPaD treatment restores memory deficits in 5xFAD mice. (A) Schedule of behavioral tests. Hes5 promoter-driven control or iPaD lentivirus was injected into the hippocampal dentate gyrus of 5xFAD mice. At 4 wpi, these mice underwent behavioral tests. Schematic was created using BioRender. (B) Traces of control and iPaD lentivirus-injected mice in the Barnes maze test. (C-E) Latency time (C), distance to reach the goal hole (D), and the number of incorrect holes visited (E) were evaluated during the training session in the Barnes maze test. n.s., not significant. Control, n = 10; iPaD, n = 12. (F) A schematic diagram of the fear conditioning test and pattern separation test. Control, n = 10; iPaD, n = 9. (G) The percentage of time spent freezing in context A was measured during days 1–3 in the fear conditioning test. (H,I) The percentage of time spent freezing in context A or C was measured during days 4–11 (blocks 1–4). (J) The percentage of time spent freezing in context A or C in block 4 is shown. Values are presented as the mean ± SEM. *P < 0.05, **P < 0.01, ***P < 0.001. Statistical significance was assessed using two-way ANOVA followed by Tukey’s post hoc test for multiple group comparisons.

We next performed a contextual fear conditioning test (Fig. 3A), where each mouse was placed in a chamber (context A) and received an electric shock after 3 min. This fear conditioning was repeated for 3 days (Fig. 3F), and on days 2 and 3, 5xFAD mice injected with the iPaD lentivirus showed longer freezing behavior than those injected with the control lentivirus (Fig. 3G), indicating that their performance was improved by iPaD treatment. After this fear conditioning test, each mouse was placed in a chamber with an electric shock (context A) and in a different chamber without an electric shock (context C) on each day (Fig. 3F). 5xFAD mice injected with the iPaD lentivirus showed a tendency of shorter freezing behavior in context C than in context A (Fig. 3I,J), whereas those injected with the control lentivirus exhibited freezing behavior for a similar amount of time in both contexts (Fig. 3H,J). These data indicate that injection of the iPaD lentivirus, which activates neurogenesis and inhibits Aβ deposition in 5xFAD mice, improves their learning and memory ability.

### Downstream events of iPaD treatment in 5xFAD mice

To understand the mechanism by which activated neurogenesis improved Aβ pathology in a non-cell-autonomous manner in the hippocampal dentate gyrus, we injected control or iPaD lentivirus into the hippocampus of 5–6-month-old 5xFAD mice, and 4 weeks later, isolated the hippocampal dentate gyrus and subjected it to RNA-seq analysis (Fig. 4A). At this stage, 5xFAD mice exhibited decreased neurogenesis and increased Aβ deposition, whereas iPaD treatment activated neurogenesis and reduced Aβ deposition (Figs. 1 and 2). We identified 592 genes that were significantly upregulated and 462 genes that were significantly downregulated by iPaD treatment in the hippocampal dentate gyrus of 5xFAD mice (Fig. 4B), suggesting that the transcriptome of this region was considerably altered by the robust activation of neurogenesis.

**Figure 4.**
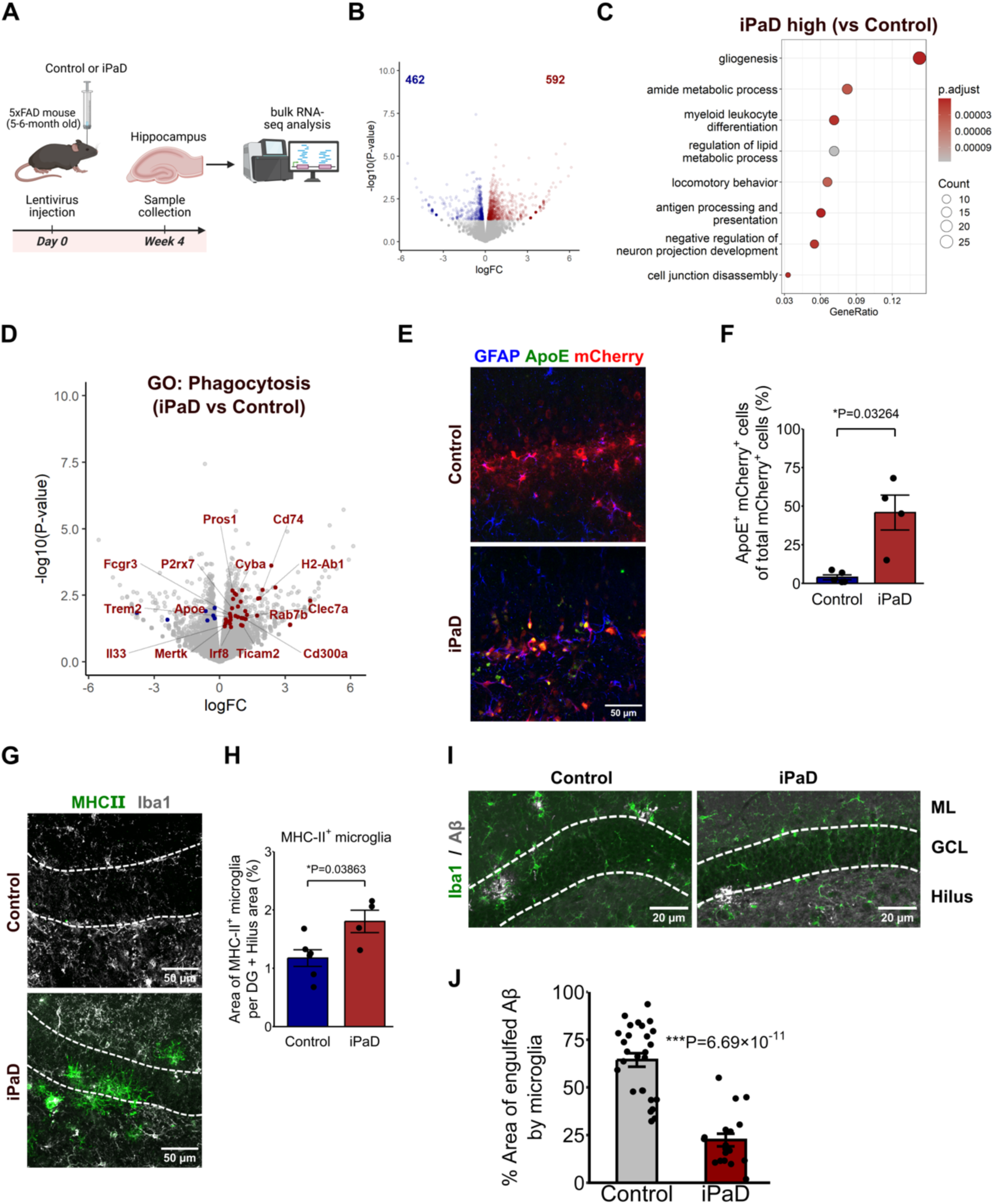
iPaD treatment activates astrocytes and microglia in 5xFAD mice. (A) Hes5 promoter-driven control or iPaD lentivirus was injected into the hippocampus of 5-to-6-month-old 5xFAD mice, and 4 weeks later the whole hippocampal dentate gyrus was subjected to RNA-seq analysis. Schematic was created using BioRender. (B) Volcano plot displaying differentially expressed genes (DEGs) in the hippocampal dentate gyrus between control and iPaD lentivirus-injected 5xFAD mice. (C) GO enrichment dot plot of upregulated genes in iPaD lentivirus-injected 5xFAD mice compared to the control. (D) Volcano plot showing DEGs associated with phagocytosis (GO:0006909); *Apoe* and *Il33* were manually included in the gene set. (E,F) Representative images (E) and quantification (F) of immunostaining for GFAP, ApoE, and mCherry in the hippocampal dentate gyrus of control or iPaD lentivirus-injected 5xFAD mice. n = 4 per group. (G,H) Representative images (G) and quantification (H) of immunostaining for Iba1, MHCII, and Aβ in the hippocampal dentate gyrus of control or iPaD lentivirus-injected 5xFAD mice. n = 4 per group. (I,J) Representative images (I) and quantification (J) of the area of Aβ plaques engulfed by microglia in the hippocampal dentate gyrus of control or iPaD lentivirus-injected 5xFAD mice. Control, n = 27; iPaD, n = 18. Values are presented as the mean ± SEM. *P < 0.05, **P < 0.01, ***P < 0.001. Statistical significance was assessed using Welch’s t test for pairwise comparisons. Scale bars: 50 µm (E,G); 20 µm (I).

Gene Ontology (GO) enrichment analyses showed that upregulated genes included those involved in glia-related activity (Fig. 4C). Many of these genes are known to ameliorate Alzheimer’s disease pathology, including genes involved in phagocytosis such as *Apoe*, *Trem2*, *Il33*, and *Mertk* (Fig. 4D) (Huang et al. 2021; Lau et al. 2023; van Olst et al. 2025). Among them, apolipoprotein E (*Apoe*), a component of high-density lipoprotein particles that transport lipids between cells, is mainly expressed by astrocytes and regulates Aβ clearance (Holtzman et al. 1999). In iPaD-treated 5xFAD mice, Apoe^+^;GFAP^+^ astrocytes were significantly increased in number compared to the control (Fig. 4E,F). In addition, interleukin-33 (*Il33*) was previously shown to induce major histocompatibility complex class II (MHC-II)^+^ microglial cells, which exhibit Aβ clearance activity (Lau et al. 2020). In iPaD-treated 5xFAD mice, MHC-II^+^;Iba1^+^ microglial cells were significantly increased in number compared to the control (Fig. 4G,H). However, more microglial cells accumulated near Aβ deposits in control mice than in iPaD mice (Fig. 4I,J), suggesting that even though microglial cells accumulate near Aβ deposits in control mice, they cannot efficiently contribute to Aβ clearance. These results suggest that the astrocytes and microglial cells activated by iPaD treatment can contribute to the non-cell-autonomous clearance of Aβ deposits, thereby ameliorating Alzheimer’s disease pathology.

Unlike the upregulated genes, the functional relevance of most of the top downregulated genes to Alzheimer’s disease is not well known. Thus, we decided to characterize these genes by examining whether knockdown of any of them could ameliorate Alzheimer’s disease pathology.

### *Prkag2, Gfra2, Rpl29*, and *Maml2* knockdown increases neurogenesis and reduces Aβ deposition

Among the top downregulated genes, we selected *Maml2*, *Ptgds*, *Prkag2*, *Gfra2*, *Rpl29*, *Itpk1*, and *Mical3*, which were expressed widely in the dentate gyrus and most significantly repressed by iPaD treatment (Fig. 5A), and examined the effects of their knockdown on Aβ pathology. Using a lentivirus, we delivered a construct of miR-E backbone short hairpin RNA (Fellmann et al., 2013) for each gene together with mCherry under the control of the elongation factor promoter, which is a ubiquitous promoter (Fig. 5B). As a control, a scrambled sequence was used (Fig. 5B). Each lentivirus was injected into the hippocampus of 5- to 6-month-old 5xFAD mice, and the effect was examined 8 weeks later (Fig. 5C). Among the tested genes, knockdown of *Prkag2*, which encodes protein kinase AMP-activated non-catalytic subunit gamma 2, most effectively increased the number of DCX^+^ neuroblasts (Fig. 5D,F) and reduced Aβ deposition (Fig. 5E,G). Knockdown of *Gfra2*, which encodes glial cell line-derived neurotrophic factor family receptor alpha 2, *Rpl29*, which encodes ribosomal protein L29, and *Maml2*, which encodes a co-activator of Notch signaling, also increased the number of DCX^+^ neuroblasts (Fig. 5D,F) and reduced Aβ deposition (Fig. 5E,G), but less effectively than *Prkag2* knockdown. We confirmed that these shRNAs effectively reduced target gene expression in cultured NSCs (Supplemental Fig. S3A-D). Notably, *Prkag2* and *Gfra2* knockdown also downregulated Aβ levels in NSCs (Supplemental Fig. S3E). These results suggest that *Prkag2, Gfra2*, *Rpl29*, and *Maml2* knockdown effectively activates neurogenesis and ameliorates Aβ pathology, suggesting that these genes could be new therapeutic targets for Alzheimer’s disease.

**Figure 5.**
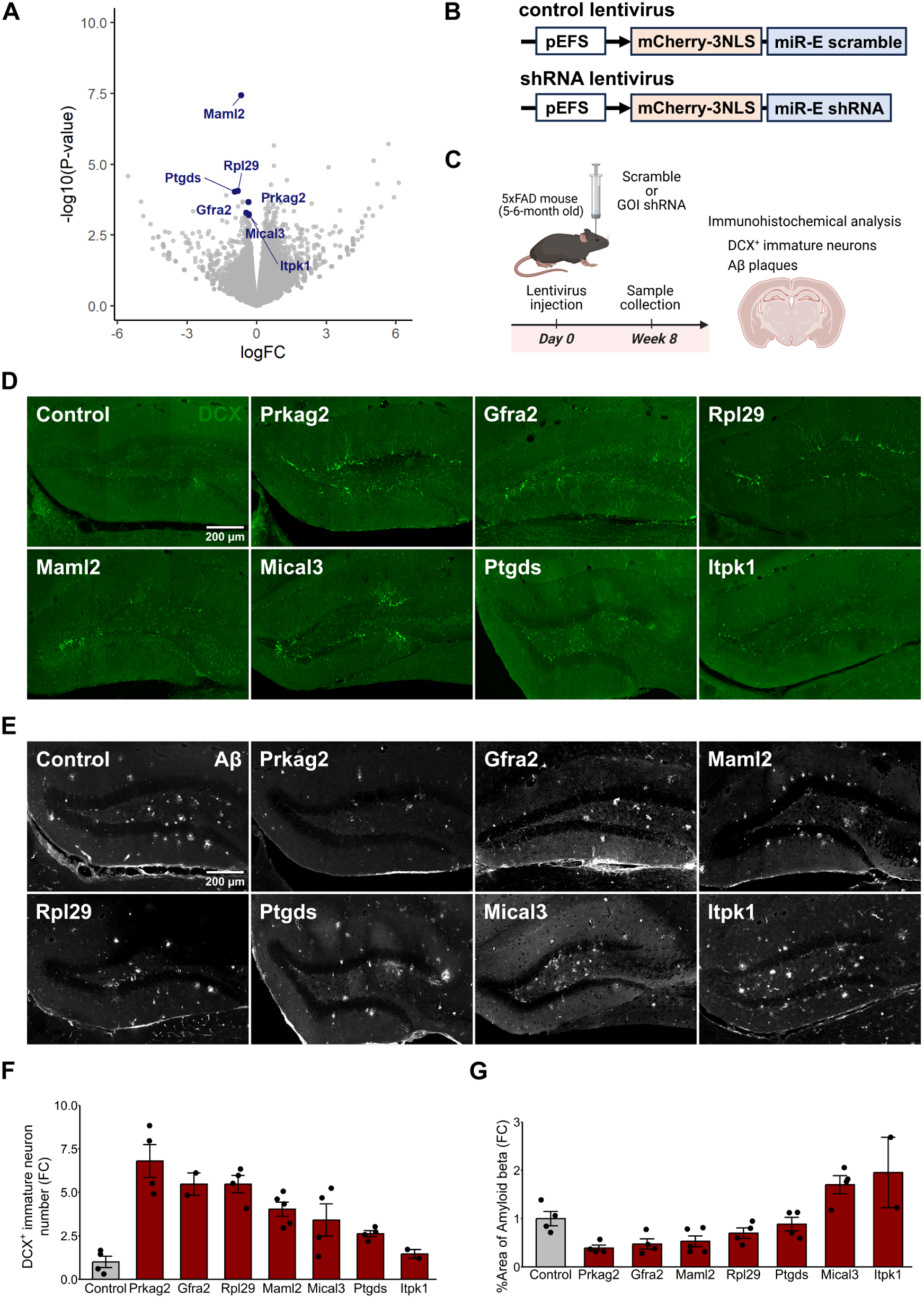
Analysis of representative genes downregulated by iPaD treatment. (A) Volcano plot showing seven representative downregulated genes with relatively high expression levels, selected from among the top-ranked candidates in the bulk RNA-seq analysis of hippocampal tissue from 5- to 6-month-old 5xFAD mice at 4 wpi with control or iPaD lentivirus. (B) Schematic structures of the elongation factor promoter (pEFS)-driven control and knockdown (shRNA) lentiviruses. (C) The schedule for lentivirus infection and immunohistochemical analysis. Schematic was created using BioRender. (D,E) Immunohistochemical analyses of DCX^+^ cells (D) and Aβ deposition (E) after pEFS-driven control or knockdown lentivirus infection. Scale bars: 200 µm. (F,G) Quantification of DCX^+^ cells (F) and Aβ deposition (G) after pEFS-driven control or knockdown lentivirus infection. Control, *n* = 4; *Prkag2*, *n* = 4; *Gfra2*, *n* = 4; *Rpl29*, *n* = 4; *Maml2*, *n* = 5; *Mical3*, *n* = 4; *Ptgds*, *n* = 4; *Itpk1*, *n* = 2. Data are presented as the mean ± SEM.

We next analyzed the expression patterns of *Prkag2, Gfra2*, *Rpl29*, and *Maml2* using a previously published single-cell transcriptome data set for the hippocampal regions of wild-type and 5xFAD mice (Habib et al., 2020). *Prkag2* was expressed by many granule cells and pyramidal neurons and subsets of glial cells in the CA regions of the hippocampus (Supplemental Fig. S4). *Maml2* was expressed mainly by granule cells, oligodendrocyte precursor cells (OPCs), and astrocytes (Supplemental Fig. S4). *Gfra2* and *Rpl29* were expressed at very low levels, but widely, in this region (Supplemental Fig. S4). These results indicated that the knockdown of these genes could affect many cell types in the hippocampus.

### iPaD treatment and *Prkag2* knockdown activate AMPK signaling

Since the knockdown of *Prkag2*, which encodes a regulatory subunit of AMPK, most effectively improved Aβ pathology, we next analyzed whether AMPK signaling is activated or inactivated by iPaD treatment and *Prkag2* knockdown. Notably, phosphorylation of the AMPK catalytic subunit PRKAA1/2 at Thr172 (pAMPK), an active form of AMPK, was increased in NSCs in 5xFAD mice by iPaD treatment (Fig. 6A,B). *Prkag2* knockdown also increased pAMPK in virus-infected cells (Fig. 6E,F). Thus, both iPaD treatment and *Prkag2* knockdown activate AMPK signaling in 5xFAD mice.

**Figure 6.**
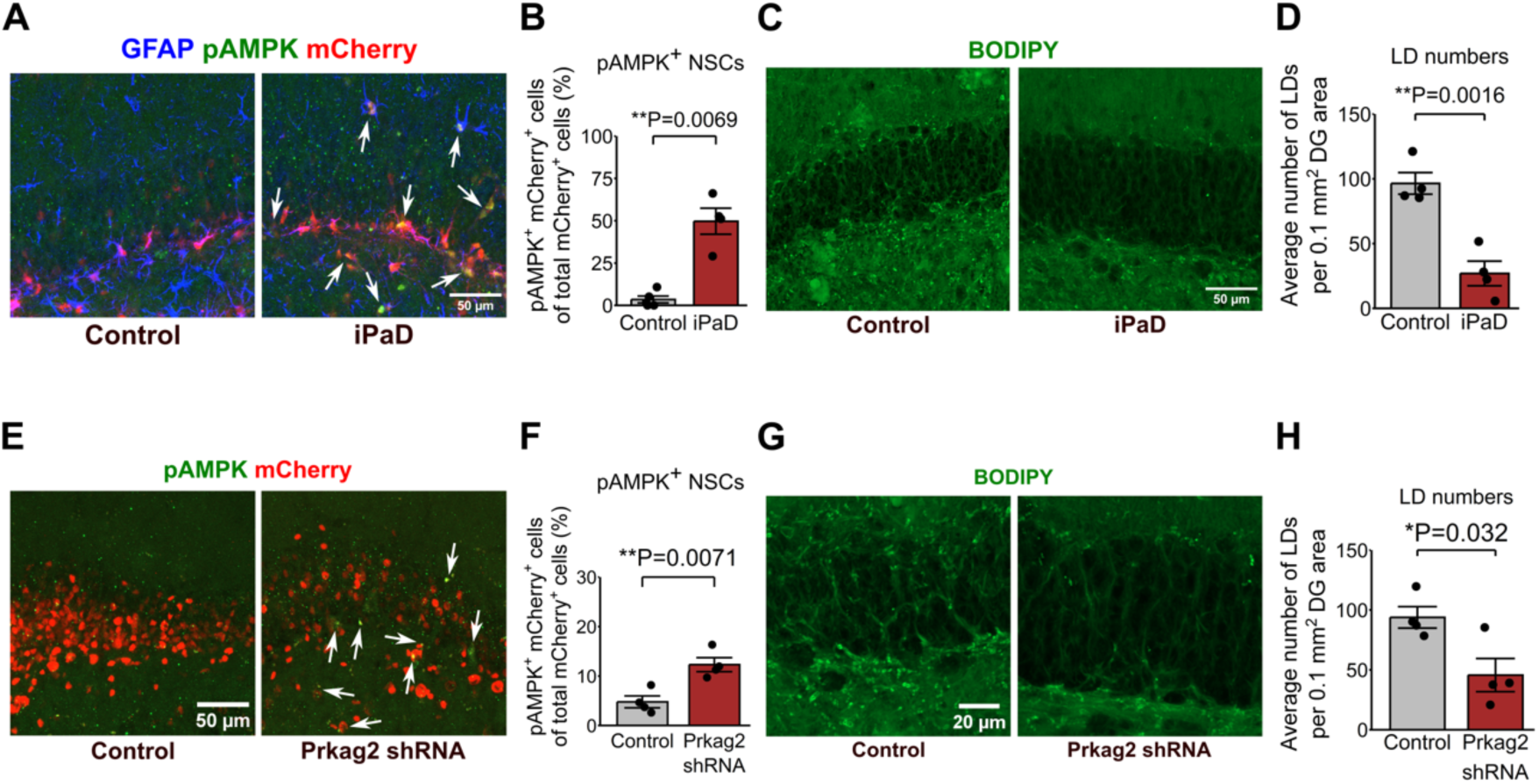
iPaD treatment and *Prkag2* knockdown activated AMPK signaling and lipid metabolism. (A,B) Immunochemical analyses (A) and quantification (B) of GFAP, pAMPK, and mCherry in the hippocampal dentate gyrus of control or iPaD lentivirus-injected 5xFAD mice. Arrows indicate GFAP^+^;pAMPK^+^;mCherry^+^ cells. (C,D) BODIPY analyses and quantification of the hippocampal dentate gyrus of control or iPaD lentivirus-injected 5xFAD mice. Control, n = 4; iPaD, n = 4. (E,F) Immunochemical analyses and quantification of pAMPK and mCherry in the hippocampal dentate gyrus of control or *Prkag2* knockdown lentivirus-injected 5xFAD mice. Arrows indicate pAMPK^+^;mCherry^+^ cells. (G,H) BODIPY analyses and quantification of the hippocampal dentate gyrus of control or *Prkag2* knockdown lentivirus-injected 5xFAD mice. Scramble control, n = 4; *Prkag2* shRNA, n = 4. Data are presented as mean ± SEM. *P < 0.05, **P < 0.01, ***P < 0.001. Statistical significance was assessed using Welch’s t test for pairwise comparisons. Scale bars: 50 µm (A,C,E); 20 µm (G).

Since AMPK signaling reduces lipid droplet accumulation, which causes pathological phenomena in the brain of patients with neurodegenerative disorders (Liu et al. 2015; Li et al. 2024), we next examined whether lipid droplet accumulation was affected by iPaD treatment and *Prkag2* knockdown. Imaging with BODIPY, a fluorescent lipid droplet indicator, showed that lipid droplets accumulated with aging in the hippocampus of 5xFAD mice whereas iPaD treatment significantly reduced lipid droplet accumulation in this region (Fig. 6C,D). Similarly, *Prkag2* knockdown effectively reduced lipid droplet accumulation in 5xFAD mice (Fig. 6G,H). These results suggest that both NSC-specific treatment with iPaD and broad knockdown of *Prkag2* in the hippocampus improve lipid metabolism in 5xFAD mice through the activation of AMPK signaling.

We next examined how *Prkag2* knockdown activates AMPK signaling. *Prkag2* knockdown in cultured NSCs increased the expression levels of the other genes encoding AMPK subunits, i.e., *Prkaa1*, *Prkaa2*, *Prkab1*, *Prkab2, Prkag1,* and *Prkag3*, (Supplemental Fig. 5A-F) as well as pAMPK (Supplemental Fig. S5G). These results suggest that *Prkag2* knockdown increases the expression levels of other AMPK subunits, leading to the activation of AMPK signaling.

*Prkag2* knockdown also activated neurogenesis in 5xFAD mice, and we next examined whether this treatment can directly activate cultured NSCs. NSCs were kept quiescent with bFGF and BMP, and after they were infected with either *Prkag2* knockdown or control lentivirus, cell proliferation was examined by EdU uptake. The proportion of EdU^+^ NSCs was increased by *Prkag2* knockdown compared to the control (Supplemental Fig. S5H). Thus, AMPK activation by *Prkag2* knockdown directly activates quiescent NSCs.

### Common downstream events of iPaD treatment and *Prkag2* knockdown in 5xFAD mice

Since *Prkag2* knockdown led to the activation of neurogenesis and inhibition of Aβ accumulation, as observed with iPaD treatment, we next examined which genes are commonly regulated by NSC-specific iPaD treatment and broad knockdown of *Prkag2*. To this end, we injected the *Prkag2* knockdown lentivirus into the hippocampus of 5- to 6-month-old 5xFAD mice, and 4 weeks later, the hippocampus was isolated and subjected to RNA-seq analysis (Supplemental Fig. S6A). We identified 3,615 genes that were significantly upregulated and 2,134 genes that were significantly downregulated in the hippocampus of 5xFAD mice by *Prkag2* knockdown (Supplemental Fig. S6B). GO enrichment analyses indicated that immune-response genes such as genes involved in lymphocyte mediated immunity, leukocyte cell-cell adhesion, and phagocytosis were upregulated by *Prkag2* knockdown (Supplemental Fig. S6C). Comparison of the transcriptomes of NSC-specific iPaD treatment and *Prkag2* knockdown revealed that immune-response genes were concomitantly upregulated (Supplemental Fig. S6D). In addition, genes associated with mitochondrial function and homeostasis were upregulated by both treatments compared to the control (Fig. 7A), indicating potential enhancement of mitochondrial activity. This effect may be due to the activation of AMPK signaling, which is known to regulate mitochondrial dynamics and quality control mechanisms such as mitochondrial biogenesis, mitophagy, oxidative phosphorylation, and ATP metabolism (Mihaylova and Shaw 2011; Herzig and Shaw 2018). Given that mitochondrial dysfunction is a hallmark of Alzheimer’s disease (Mary et al. 2022), these findings suggest that iPaD treatment and *Prkag2* knockdown may restore mitochondrial health and bioenergetic capacity. In parallel, genes involved in lipid catabolism, fatty acid metabolism/biosynthesis, and carboxylic acid metabolism/biosynthesis were upregulated by iPaD treatment and *Prkag2* knockdown, whereas genes related to mRNA metabolism were downregulated (Fig. 7B,C). Notably, *Apoe*, a gene involved in lipid processing and Aβ clearance (Koistinaho et al., 2004; Jiang 2008), was commonly upregulated by iPaD treatment and *Prkag2* knockdown (Fig. 7B), suggesting a functional link between lipid metabolism and amyloid degradation. Furthermore, genes involved in autophagy and microglial activation were also commonly upregulated by iPaD treatment and *Prkag2* knockdown (Fig. 7D,E), potentially contributing to Aβ clearance. In contrast, genes involved in senescence were commonly downregulated (Fig. 7G). Together, these results support the notion that both iPaD treatment and *Prkag2* knockdown stimulate AMPK signaling, thereby improving mitochondrial function, metabolic homeostasis, and cellular clearance mechanisms in the Alzheimer’s disease brain.

**Figure 7.**
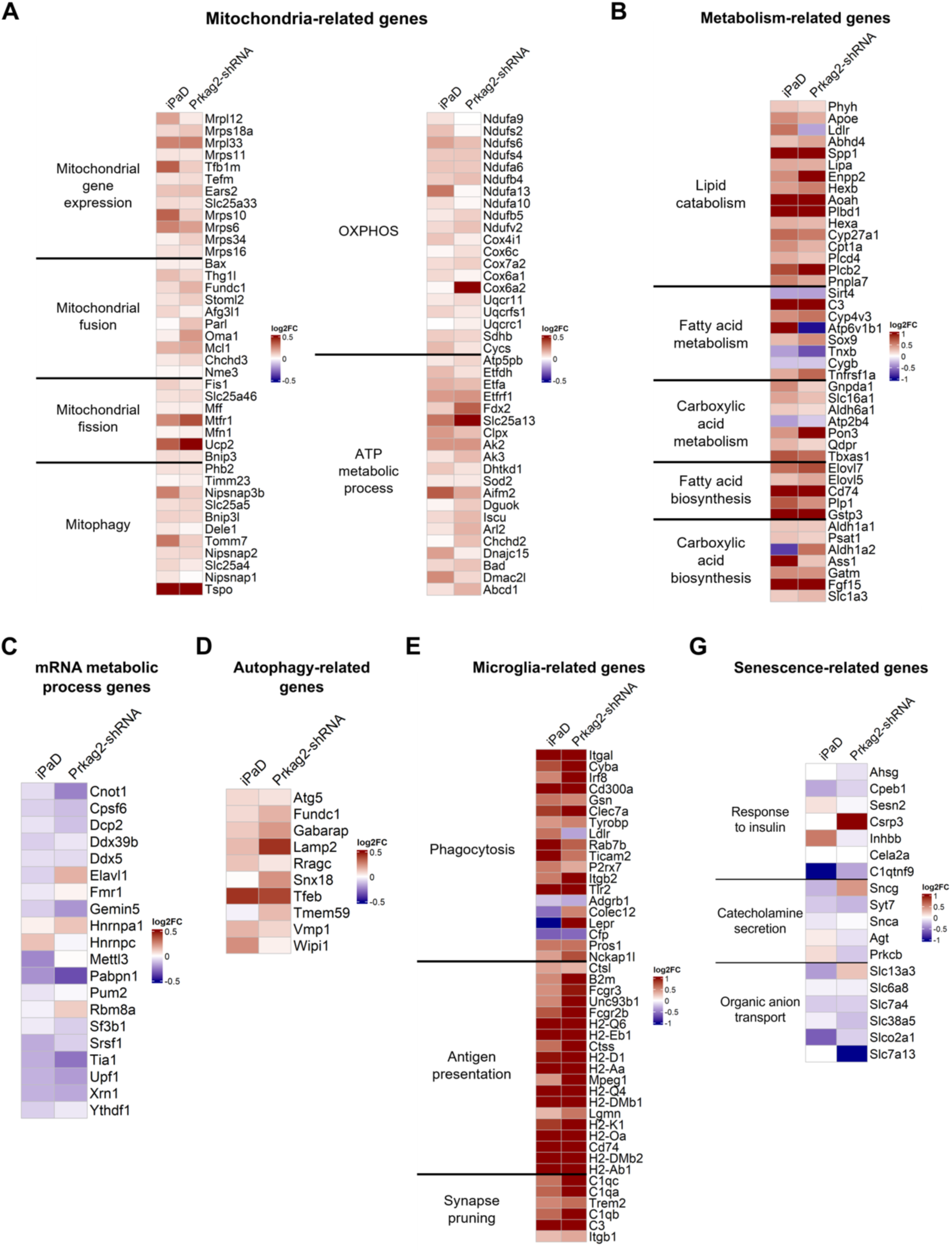
Common downstream genes of iPaD treatment and *Prkag2* knockdown in 5xFAD mice. Bulk RNA-seq analyses of the hippocampal dentate gyrus from 5–6-month-old 5xFAD mice at 4 wpi with control or iPaD lentivirus, or with scramble control or Prkag2 shRNA lentivirus. (A) Representative heatmaps of mitochondria-related genes. (B) Representative heatmaps of metabolism-related genes. (C) Representative heatmaps of mRNA metabolism-related genes (GO:0016071, mRNA metabolic process). (D) Representative heatmaps of autophagy-related genes (GO:0006914, autophagy). (E) Representative heatmaps of microglia-related genes associated with phagocytosis (GO:0006909), antigen presentation (GO:0048002), and synapse pruning (GO:0098883). (F) Representative heatmaps of senescence-related genes, including those involved in the response to insulin (GO:0032868), catecholamine secretion (GO:0050432), and organic anion transport (GO:0015711), selected using SenePy.

Analysis of the published single-cell RNA-seq data (Habib et al. 2020) indicated that *Prkag2* expression, which was widely observed in many cell types, was upregulated in 5xFAD and aged brains (Supplemental Fig. S7A-C). GO enrichment analyses indicated that in *Prkag2*-expressing cells, genes involved in cytoplasmic translation and oxidative phosphorylation as well as mitochondria-related genes were downregulated compared to *Prkag2*-negative cells (Supplemental Fig. S7D,E), suggesting that *Prkag2* upregulation may lead to cellular dysfunction. Thus, these results suggest that *Prkag2* knockdown normalizes cellular function by upregulating the expression of various genes including many mitochondria-related genes in 5xFAD mice.

## Discussion

### iPaD treatment improves Aβ pathology and cognitive function in Alzheimer’s disease model mice

The activation of adult neurogenesis has been linked to several neurodegenerative disorders, leaving unanswered questions about their potential therapeutic benefits. Previous studies reported that while exercise can reduce Aβ deposition, the activation of adult neurogenesis alone is insufficient to achieve this effect in Alzheimer’s disease model mice (Choi et al., 2018; Li et al., 2023). However, our findings demonstrate that the activation of adult neurogenesis by iPaD treatment is sufficient to reduce Aβ deposition and enhance cognitive function. iPaD treatment induces adult neurogenesis with markedly higher efficiency (∼4-fold) than other methods (∼1.5-fold; Choi et al., 2018; Li et al., 2023), suggesting that this elevated level of neurogenesis may yield distinct effects on Aβ pathology. Only a small subset of newly-born neurons integrate into hippocampal neural circuits, while the majority undergo apoptosis and are subsequently phagocytosed by microglia (Sierra et al. 2010). Thus, iPaD treatment likely activates a greater number of microglial cells than previous methods, thereby enhancing Aβ clearance through microglial phagocytosis. Indeed, we observed the upregulation of microglia-associated clearance factors such as compliment factor C3 and Il33 (Maier et al. 2008; Lau et al. 2020) following iPaD treatment. Additionally, MHC-II^+^ microglial cells, which exhibit Aβ clearance activity (Lau et al. 2020), were significantly increased in number. iPaD treatment also reduced lipid droplet accumulation, which is associated with enhanced phagocytic activity, further supporting increased Aβ clearance (Lau et al. 2023; Wu et al. 2025). Beyond microglia, astrocytes also contribute to Aβ clearance. Apoe, which is secreted by astrocytes, plays a critical role in Aβ metabolism (Holtzman et al. 1999). We found that the number of Apoe^+^ astrocytes was increased by iPaD treatment, suggesting a cooperative role of astrocytes and microglia in mediating Aβ clearance.

In addition to these cellular clearance mechanisms, transcriptomic analysis revealed that iPaD treatment upregulates genes associated with mitochondrial function and metabolic homeostasis. Mitochondrial dysfunction is a well-established feature of Alzheimer’s disease, contributing to impaired energy production and increased oxidative stress (Mary et al. 2022). iPaD treatment appears to restore mitochondrial gene expression and promote oxidative metabolism, which may support both neurogenesis and microglial function. This is consistent with known roles of AMPK signaling in regulating mitochondrial homeostasis and lipid catabolism (Mihaylova and Shaw 2011; Herzig and Shaw 2018). Thus, iPaD may improve the overall metabolic state of neuronal and glial cells, further enhancing their capacity to clear Aβ and support cognitive function.

Together, these findings indicate that the robust induction of adult neurogenesis by iPaD treatment not only promotes cellular mechanisms of Aβ clearance via microglia and astrocytes but also improves mitochondrial activity and energy metabolism, contributing to the amelioration of Alzheimer’s disease pathology.

### Downstream effects of *Prkag2* knockdown

Among the downstream genes of iPaD treatment, *Prkag2* emerged as one of the most promising candidates for Alzheimer’s disease therapy, as its knockdown most effectively enhanced neurogenesis and Aβ clearance. *Prkag2* encodes the γ2 regulatory subunit of AMPK, a heterotrimer composed of α (α1 and α2), β (β1 and β2), and γ (γ1, γ2, and γ3) subunits that is a key enzyme involved in cellular energy metabolism. Previous studies have reported that PRKAG2 expression is upregulated in postmortem brains of Alzheimer’s disease patients (Bharadwaj and Martins 2020). However, it was unclear whether this upregulation contributes to Aβ pathology or represents a compensatory response. Our data show that *Prkag2* knockdown significantly reduced Aβ deposition in 5xFAD mice, suggesting that suppressing *Prkag2* expression may be beneficial for Aβ pathology not only in mice but also potentially in humans. Interestingly, *Prkag2* knockdown resulted in the activation of AMPK signaling, likely through the compensatory upregulation of other AMPK subunits. Although the functional distinctions between AMPK subunit isoforms are not understood fully, all γ subunits (PRKAG1, PRKAG2, and PRKAG3) contribute to AMPK activity in the brain (Cheung et al. 2000). Notably, previous studies have shown that PRKAG1, but not PRKAG2 or PRKAG3, plays a critical role in mediating AMPK activation by metformin (An et al. 2020), suggesting functional differences among the γ subunits. Therefore, it is possible that an age-related or Alzheimer’s disease-associated shift toward higher PRKAG2 expression relative to PRKAG1 could dampen AMPK signaling. Conversely, reducing PRKAG2 expression may restore or enhance AMPK signaling by promoting the expression or activity of other subunits.

AMPK activation reduces lipid droplet accumulation and stimulates autophagy, both of which contribute to the mitigation of Alzheimer’s disease pathology (Wang and Jia 2023; Li et al., 2024). Furthermore, our data demonstrate that *Prkag2* knockdown promoted NSC proliferation, consistent with previous findings that metformin, an AMPK activator used in type 2 diabetes mellitus, enhances NSC proliferation (Wang 2012; Xing et al., 2024). Collectively, these findings suggest that *Prkag2* knockdown induces neurogenesis and alleviates Aβ pathology by activating AMPK signaling.

Recent studies have shown that metformin not only manages type 2 diabetes mellitus but also improves cognitive function and promotes healthy aging (Hsu et al. 2011). However, conflicting reports indicate that long-term metformin treatment can impair cognition in Alzheimer’s disease model mice (Cho et al., 2024). In addition, the expression of AMPKα1, a catalytic subunit of AMPK, is elevated in the brain of Alzheimer’s disease patients, and its repression alleviates cognitive deficits in Alzheimer’s disease model mice (Zimmermann et al. 2020), although it remains unclear whether this repression leads to overall AMPK activation or inhibition. These findings underscore the need for further investigation into the long-term effects of AMPK signaling, particularly in the context of Alzheimer’s disease.

While our study focused on Alzheimer’s disease model mice, AMPK activation is also recognized as beneficial in other neurological disorders, including Parkinson’s disease and Huntington’s disease (Vázquez-Manrique et al. 2016; Curry et al. 2018; Ripa et al. 2023). Thus, *Prkag2* knockdown may represent a broadly applicable therapeutic strategy for treating various neurodegenerative diseases and promoting healthy longevity.

### Downstream effects of *Gfra2, Rpl29,* and *Maml2* knockdown

*Gfra2*, *Rpl29*, or *Maml2* knockdown effectively enhanced adult neurogenesis and promoted Aβ clearance as well, indicating that these genes are also promising therapeutic targets for Alzheimer’s disease. GFRA2 expression is elevated in postmortem brains of Alzheimer’s disease patients (Haytural et al. 2021), although its specific role in the disease remains unclear. Its ligand, glial cell line-derived neurotrophic factor (GDNF), regulates axonal growth and neuronal survival; however, neurons from Alzheimer’s disease patients exhibit a diminished response to GDNF (Konishi et al. 2014). Notably, GDNF levels are significantly reduced in the serum, but are elevated in the cerebrospinal fluid of Alzheimer’s disease patients compared to age-matched controls (Straten et al. 2009). This suggests that the increase of GDNF in cerebrospinal fluid may represent a compensatory mechanism to enhance neurotrophic support (Straten et al. 2009). Since GDNF binds preferentially to a different receptor subtype, GFRA1, knockdown of *Gfra2* may redirect GDNF signaling through GFRA1, potentially restoring its neuroprotective effects. However, *Gfra2*-null mice exhibit behavioral impairments and memory deficits (Võikar et al. 2004), indicating that a certain level of GFRA2 is necessary for normal brain function.

*Rpl29* encodes a ribosomal protein, although its function in Alzheimer’s disease is not known. The altered expression of many *Rpl* genes has been observed in Alzheimer’s disease patients, implicating dysregulated protein synthesis in disease pathology (Hernández-Ortega et al. 2015). Therefore, *Rpl29* knockdown may help normalize translational control, although the underlying mechanisms require further investigation.

MAML proteins are a family of three co-activators for various transcription factors such as the Notch signaling factor RBPj (Lin et al. 2002; Wu et al., 2002) and myocyte enhancer factor 2C (MEF2C) (Shen et al. 2006). Thus, *Maml2* knockdown likely leads to the broad downregulation of gene expression, including genes involved in Notch signaling. We found that *Maml2* knockdown significantly enhanced neurogenesis and reduced Aβ deposition. In the adult brain, NSC quiescence is maintained by the high expression of the Notch effector HES1 (Sueda et al. 2019). Thus, *Maml2* knockdown may reduce HES1 expression, resulting in the activation of quiescent NSCs. Moreover, recent reports indicate that *Maml2* is upregulated in the brains of Alzheimer’s disease patients (Xiong et al. 2023), supporting the potential therapeutic relevance of its knockdown in both animal models and humans.

*Prkag2, Gfra2, Rpl29,* or *Maml2* knockdown activated adult neurogenesis and reduced Aβ deposition. To our knowledge, this is the first report showing that knockdown of a single gene can both activate adult neurogenesis and reduce amyloid-β pathology in Alzheimer’s disease model mice. While these genes displayed similar beneficial effects on Alzheimer’s disease pathology, it remains unknown whether they function in the same pathway or independently. However, we observed a strong correlation between the extent of neurogenesis activation and the degree of Aβ reduction, suggesting a reciprocal relationship between these two processes. Indeed, Aβ oligomers are known to inhibit adult neurogenesis in Alzheimer’s disease animal models (Scopa et al. 2020), suggesting that reducing the Aβ burden may help restore neurogenic capacity. Conversely, enhancing neurogenesis may delay the progression of neurodegenerative diseases such as Alzheimer’s disease and Huntington’s disease (Benraiss et al. 2013; Choi et al. 2018). Collectively, these findings support the idea that the targeted knockdown of these genes, particularly *Prkag2,* is a promising therapeutic strategy not only for Alzheimer’s disease but also for a broad range of neurodegenerative disorders.

## Materials and Methods

### Animals

5xFAD mice (Oakley et al. 2006) were obtained from The Jackson Laboratory and used as Alzheimer’s disease model mice. In brief, 5xFAD mice carry doubly transgenic constructs of mutant human amyloid precursor protein (APP) (the Swedish mutation: K670N, M671L; the Florida mutation: I716V; the London mutation: V717I) and presenilin 1 (PS1) (M146L; L286V) under the control of the mouse Thy1 promoter (Tg6799 line). The 5×FAD lines (B6/SJL genetic background) were procured from Jackson Laboratory and backcrossed for more than 2 generations with C57BL/6J wildtype (WT) mice. WT littermate mice were used as control animals. All mice were handled following the guidelines outlined in the Kyoto University and RIKEN Guide for the Care and Use of Laboratory Animals. The experimental protocols received approval from the Experimental Animal Committee of the Institute for Frontier Life and Medical Sciences, Kyoto University, and the Animal Experiment Committee at RIKEN.

### Construction of lentiviruses

The sequence for each gene was subcloned into the CSII-1.6-kb-pmHes5-mCherry-3NLS-MCS plasmid for lentivirus production. For lentivirus production, 134.4 μg of each CSII plasmid, 99.2 μg of gag-pol plasmid (psPAX2), and 30.72 μg of VSV-G envelope plasmid (pMD2.G) were co-transfected into HEK293 cells plated in eight 15-cm dishes using 1.376 ml of 1 mg/ml Polyethylenimine-MAX (Polysciences). The cells were cultured in OptiPRO serum-free medium (Gibco) containing 10μM Forskolin, and the medium was harvested 3 days later. A total of 160 ml of the medium was concentrated into 100 μl of PBS through centrifugation at 8,000g for 12 hours and 13,000g for 4 hours. The lentivirus titer was measured by infecting cultured NSCs, typically yielding a titer of 2 × 10^7^ units/ml. Lentiviral vectors with the miR-E backbone were used for gene expression knockdown. The following sequences were used for knockdown:

*Dyrk1a*: TTTAATTACAAAATCAATATGG

*Maml2*: TTTATCTACTAGATGACAGTCT

*Rpl29*: TTCGTATCTTTGCGACCGGGGT

*Ptgds*: TCTTGTTGAAAGTTGGGCTGCA

*Prkag2*: TTTATGAAATCTGTAATCGTGA

*Gfra2*: TTGATCACCTTTTTACTCCCTG

*Itpk1*: TTAATGATGATGTCAATTCCAA

*Mical3*: TTTTCAATAACAACCACCTTGG

As a negative control, scramble sequences were used.

### Lentivirus infection in mice

For in vivo screening and neurogenesis induction, 1 μL of lentivirus (3.75 × 10^5^ U/mL) was stereotactically delivered into the dentate gyrus bilaterally with the following coordinates: anteroposterior = −2.0 mm from bregma, lateral = ±1.4 mm, and ventral = −2.0 mm, at a flow rate of 0.3 μL/min. Brain sections were subsequently analyzed by immunohistochemistry.

### Immunohistochemistry and BODIPY staining

For the immunohistochemical analysis of brain tissues, mice underwent transcardial perfusion with 4% paraformaldehyde in PBS. Brains were subsequently postfixed with 4% paraformaldehyde in PBS, cryoprotected through sequential overnight incubation with 30% sucrose in PBS twice, embedded, and frozen in OCT (Tissue TEK). Serial 50-μm thick coronal frozen sections were obtained using a cryostat and stored at -80°C in cryoprotectant solution (BIE, 006799) until use. The following primary antibodies (final dilution and source) were used: rabbit anti-doublecortin (1:500; CST 14802), rabbit anti-MCM2 (1:500; Abcam ab4461), rat anti-mCherry (1:500; Invitrogen M11217), and mouse anti-β-amyloid (Clone 6E10; 1:500; BioLegend 9320). The first antibodies were applied for 2 overnight periods, and the second antibodies were applied for 1 overnight period. Sections were mounted with Fluoromount-G (SBA, 0100-01).

For BODIPY staining, brains were postfixed with 4% paraformaldehyde in PBS for 48 hours after perfusion fixation, embedded, and frozen in OCT. Serial 50-μm thick coronal frozen sections were stored at -20 ℃ in cryoprotectant solution (BIE, 006799) until use. Free-floating sections were washed three times in PBS followed by 1 hour blocking in PBS with 10% donkey serum. Sections were incubated in PBS with 10% donkey serum and primary antibodies for 48 hours at 4°C. After primary antibody incubation, sections were washed three times in PBS and incubated in PBS with 10% donkey serum and second antibodies for 3 hours at room temperature. Sections were washed once and incubated in PBS with BODIPY 493/503 (1:1000 from a 1 mg/mL stock solution in DMSO) for 15 min at room temperature. Sections were mounted with Fluoromount-G (SBA, 0100-01).

### Quantification of labeled cells and amyloid-β plaques

More than three mice were analyzed for each experiment. Every 7th section was evaluated for each mouse. Immunostaining was applied to all sections, except for the damaged ones, for labeling. Subsequently, z-stack images were captured using confocal microscopy (LSM780 or LSM980) with 10x or 20x objectives. Stained cells were quantified using ImageJ Fiji and expressed as the number of cells per mm^3^ for each image.

A series of every 7th brain section from each mouse was labeled with a mouse anti-amyloid-β primary antibody (Clone 6E10; 1:500; BioLegend 9320) and detected by a fluorescently labeled secondary antibody. Following immunostaining, z-stack images were obtained using confocal microscopy (LSM780 or LSM980) with 10x or 20x objectives. The number of Aβ plaques was quantified using ImageJ Fiji after adequate thresholding and noise despeckling. Results were expressed as the covered area of plaques per mm^3^ for each image.

### Behavior tests

8-12 male mice per group were used for each behavioral analysis. Mice were automatically tracked to detect multiple body points using the ANY-maze software (Stoelting Co.). Spatial memory was measured using Barnes circular maze (92 cm in diameter, 12 holes, Brain Science Idea, Kyoto, Japan), as previously described (Kaise et al., 2022). The test area was enclosed by a black curtain, with four distinctive cues placed at least 30 cm from the maze’s edge. The light intensity was set at 350 lux. The maze was cleaned with 70% ethanol and dried between each animal’s use. For habituation, each mouse was allowed to explore the maze with one escape hole for 5 minutes. The following day, each mouse underwent two training sessions per day to learn the location of one escape hole. The escape hole position remained constant for each mouse during tests but varied between mice. At the start of training, mice were placed for 30 seconds in a white cylinder at the center of the maze. The cylinder was removed as the recording began. Recordings were automatically stopped after a maximum of 5 minutes or when the mouse entered the escape hole or hovered around it for 10 seconds. The latency, distance, and errors to enter the goal hole were measured by ANY-maze software.

Contextual memory was evaluated using a contextual fear conditioning test, modified from a previously described protocol (Nakashiba et al. 2012; Walgrave et al. 2021). A conditioning chamber (17 × 10 × 10 cm), with a floor of 24 stainless steel rods connected to a shock generator, was placed inside a sound-attenuating cubicle with 60 dB background white noise. On day 1, each mouse was placed in context A (transparent walls, white light, handled with white latex gloves and white experimental clothes) for 3 minutes, followed by a 2-second, 80 mA foot shock. Mice remained in the chamber for another minute. The chamber and rods were cleaned with 70% ethanol between each use. After 24 and 48 hours, mice were reintroduced to context A, and the procedure from day 1 was repeated. The first 3 minutes of freezing behavior were measured using TimeFZ2 (O’HARA & CO., LTD, Japan).

Pattern separation was assessed using an adapted contextual fear conditioning protocol. Over 8 days (Days 4–11), freezing behavior in contexts A and C was recorded. On day 4, mice were placed in context A for 3 minutes, followed by a 2-second, 80 mA foot shock. They were allowed to remain in the chamber for an additional minute and then transferred to their housing cage. Approximately 120–150 minutes later, they were placed in context C (no foot shock, white and black vertical striped walls, blue light, handled with blue nitrile gloves and blue experimental clothes) for 4 minutes. On day 5, the context presentation order was reversed, starting with context C. From days 6-11, the procedures from days 4-5 were repeated. The first 3 minutes of freezing behavior were measured using TimeFZ2 (O’HARA & CO., LTD, Japan).

### Preparation of hippocampal dentate gyrus tissues from lentivirus-injected 5xFAD mice and transcriptome analysis

Four weeks after stereotactic injection of lentiviruses into the hippocampus of 5-to-6-month-old 5xFAD mice, total RNA was extracted from the dentate gyrus tissues using the RNeasy Mini Kit (Qiagen). The quality of RNA was assessed using an RNA Bioanalyzer, and all samples confirmed an RNA Integrity Number (RIN) above 9. Sequencing libraries were prepared with the NEBNext Ultra II Directional RNA Library Prep Kit for Illumina (NEB). Paired-end sequencing was conducted using HiSeq X (Illumina). cDNA sequences were aligned to the mouse reference genome mm10 and counted using STAR, with counts normalized using edgeR. Samples with outliers were excluded from the analyses. Differential gene expression analysis and GO enrichment analysis were performed using edgeR and clusterProfiler packages in R. For shRNA knockdown screening, genes with logCPM > 5 were retained for further analysis. Heatmaps were generated based on log fold changes (logFC) of normalized counts using the pheatmap and ComplexHeatmap R package. Mitochondria-related genes were obtained from the MitoCarta database (Rath et al., 2021), and senescence-related genes were obtained from the SenePy database (Sanborn et al., 2025).

### Mouse neural stem cell culture

Neural stem cells (NSCs) were cultured in growth medium consisting of 20 ng/mL EGF (Thermo, 53003-018), 20 ng/mL bFGF (Wako, 060–04543), penicillin/streptomycin (Nacalai Tesque, 09367-34), and N-2 MAX media supplement (R&D Systems, AR009) in DMEM/F-12 (Gibco, 11039-021), on dishes coated with Matrigel (1:667 dilution; Corning, 356231). To assess shRNA knockdown efficiency, NSCs were infected with lentivirus encoding shRNA one day after plating. Cells were harvested at three days post-infection.

For quantification of AMPK subunit gene expression, western blotting of phosphorylated AMPK, and EdU proliferation assay, the culture medium was replaced with quiescence-inducing medium [50 ng/mL BMP4 (R&D Systems, 314-BP), 20 ng/mL bFGF, penicillin/streptomycin, and N-2 MAX supplement in DMEM/F-12] one day after plating. NSCs were infected with shRNA lentivirus 48 hours later, and samples were collected at 72 hours post-infection.

For EdU proliferation assay, 10 µM EdU was added 48 hours after infection. After an additional 24 hours, cells were fixed with 4% paraformaldehyde, and EdU incorporation was visualized using the Click-iT™ EdU Imaging Kit (Invitrogen, C10337). Images were acquired using a BZ-X710 All-in-One Fluorescence Microscope (Keyence), and EdU⁺;mCherry⁺;DAPI⁺ cells were quantified using ImageJ Fiji.

### Quantitative RT-PCR

Total RNA was extracted from mouse hippocampal tissues and cultured NSCs using the RNeasy Mini Kit (QIAGEN, 74104) according to the manufacturer’s instructions. A total of 200–500ng of RNA was reverse-transcribed into cDNA using the ReverTra Ace® qPCR RT Kit (TOYOBO, FSQ-101). Quantitative RT-PCR was performed using the KOD SYBR qPCR Mix (TOYOBO, QKD-201) on a QuantStudio 12K Real-Time PCR System (Applied Biosystems). Expression levels of target genes were normalized to the housekeeping gene *Actb*, and relative fold changes were calculated using the 2^–ΔΔCt method.

### ELISA

Aβ40 levels in the culture supernatant of NSCs were measured using an ELISA kit (Wako, 294-64701) according to the manufacturer’s instructions. Absorbance was read at 450 nm using a multimode plate reader (Thermo Fisher Scientific).

### Western blotting

Cells were washed with cold PBS and lysed on ice for 30 min in lysis buffer containing 50 mM Tris-HCl (pH 8.0), 100 mM NaCl, 5 mM MgCl₂, 0.5% (w/v) Nonidet P-40, Complete™ protease inhibitor cocktail (Roche Diagnostics, Basel, Switzerland), 1 mM phenylmethylsulfonyl fluoride, 250 U/mL Benzonase (Sigma), 10 mM β-glycerophosphate, 1 mM sodium orthovanadate, 1 mM sodium fluoride (NaF), and 1 mM sodium pyrophosphate. Lysates were subjected to SDS-PAGE. The following primary antibodies were used: rabbit anti-β-actin (1:2000, Sigma-Aldrich, A2066) and rabbit anti-phospho-AMPKα (Thr172) (1:1000, Cell Signaling Technology, 2535). HRP-conjugated anti-rabbit IgG (1:2000, Cytiva, NA9340) was used as the secondary antibody. Western blot signals were visualized by chemiluminescence using Amersham ECL or ECL Prime detection reagents (Cytiva, RPN2232), and images were acquired using the ChemiDoc™ Touch Imaging System (Bio-Rad). Band intensities were quantified using ImageJ Fiji and normalized to the corresponding β-actin signal.

### Statistical analyses

Data are presented as means ± SEM. Statistical analyses were conducted using the R software. Statistical significance was assessed using Welch’s t test for pairwise comparisons, and one-way or two-way ANOVA followed by Tukey’s post hoc test for multiple group comparisons. P values < 0.05 were considered statistically significant.

### Single-nucleus RNA-sequencing analysis

The snRNA-seq dataset of the hippocampus of wildtype and 5xFAD mice was obtained from the GEO database accession number GSE143758 (Habib et al., 2020). The Seurat package (v5.0.1) was used to generate the count matrix and downstream analysis (Hao et al., 2023). Low-quality nuclei were filtered by the following parameters: “nCount_RNA” > 300, “nCount_RNA” < 2000, and “percent.mt” < 0.75. Dataset normalization and scaling were performed using the SCTransform function. Integration of wildtype and 5xFAD data was performed with the IntegrateData function. Clusters were then annotated manually based on the expression levels of putative markers.

## Acknowledgements

We thank Yasuharu Ishihara and Yoshikuni Tabata for critical discussion, and Kotari Ishikawa for technical help. This work was supported by Core Research for Evolutional Science and Technology (JP20gm1110002 to R.K.) from Japan Agency for Medical Research and Development, Specially Promoted Research from Japan Society for the Promotion of Science (21H04976 to R.K.), and Eisai Co., Ltd.

## Author contribution

M.F. performed the experiments, analyzed the data, and wrote the manuscript; T.K. analyzed the data; T.M. performed single-nucleus RNA-sequencing analyses; T.S. performed NSC culture experiments; R.K. designed and supervised the project and wrote the manuscript.

## References

An H, Wang Y, Qin C, Li M, Maheshwari A, He L. 2020. The importance of the AMPK gamma 1 subunit in metformin suppression of liver glucose production. Sci Rep 10: 10482.

Benraiss A, Toner MJ, Xu Q, Bruel-Jungerman E, Rogers EH, Wang F, Economides AN, Davidson BL, Kageyama R, Nedergaard M, et al. 2013. Mobilization of endogenous progenitor cells regenerates functionally-integrated medium spiny striopallidal projection neurons and delays disease progression in a transgenic model of Huntington’s disease. Cell Stem Cell 12: 787–799.

Bharadwaj P, Martins RN. 2020. PRKAG2 gene expression is elevated and its protein levels are associated with increased Amyloid-β accumulation in the Alzheimer’s disease brain. J Alzheimer’s Dis 74: 441–448.

Cheung PCF, Salt IP, Davies SP, Hardie DG, Carling D. 2000. Characterization of AMP-activated protein kinase γ-subunit isoforms and their role in AMP binding. Biochem J 346: 659–669.

Cho SY, Kim EW, Park SJ, Phillips BU, Jeong J, Kim H, Heath CJ, Kim D, Jang Y, López-Cruz L, et al. 2024. Reconsidering repurposing: long-term metformin treatment impairs cognition in Alzheimer’s model mice. Transl Psychiatry 14: 34.

Choi SH, Bylykbashi E, Chatila ZK, Lee SW, Pulli B, Clemenson GD, Kim E, Rompala A, Oram MK, Asselin C, et al. 2018. Combined adult neurogenesis and BDNF mimic exercise effects on cognition in an Alzheimer’s mouse model. Science 361: eaan8821.

Curry DW, Stutz B, Andrews ZB, Elsworth JD. 2018. Targeting AMPK signaling as a neuroprotective strategy in Parkinson’s disease. J Parkinsons Dis 8: 161–181.

Díaz-Moreno M, Armenteros T, Gradari S, Hortigüela R, García-Corzo L, FontánLozano Á, Trejo JL, Mira H. 2018. Noggin rescues age-related stem cell loss in the brain of senescent mice with neurodegenerative pathology. Proc Natl Acad Sci USA 115: 11625–11630.

Encinas JM, Michurina TV, Peunova N, Park J-H, Tordo J, Peterson DA, Fishell G, Koulakov A, Enikolopov G. 2011. Division-coupled astrocytic differentiation and age-related depletion of neural stem cells in the adult hippocampus. Cell Stem Cell 8: 566–579.

Fellmann C, Hoffmann T, Sridhar V, Hopfgartner B, Muhar M, Roth M, Lai DY, Barbosa IAM, Kwon JS, Guan Y, et al. 2013. An optimized microRNA backbone for effective single-copy RNAi. Cell Rep 5: 1704–1713.

Gonçalves JT, Schafer ST, Gage FH. 2016. Adult neurogenesis in the hippocampus: from stem cells to behavior. Cell 167: 897–914.

Habib N, McCabe C, Medina S, Varshavsky M, Kitsberg D, Dvir-Szternfeld R, Green G, Dionne D, Nguyen L, Marshall JL, et al. 2020. Disease-associated astrocytes in Alzheimer’s disease and aging. Nat Neurosci 23: 701–706.

Hao Y, Stuart T, Kowalski MH, Choudhary S, Hoffman P, Hartman A, Srivastava A, Molla G, Madad S, Fernandez-Granda C, et al. 2023. Dictionary learning for integrative, multimodal and scalable single-cell analysis. Nat Biotechnol 42: 293–304.

Haytural H, Benfeitas R, Schedin-Weiss S, Bereczki E, Rezeli M, Unwin RD, Wang X, Dammer EB, Johnson ECB, Seyfried NT, et al. 2021. Insights into the changes in the proteome of Alzheimer disease elucidated by a meta-analysis. Sci Data 8: 312.

Hernández-Ortega K, Garcia-Esparcia P, Gil L, Lucas JJ, Ferrer I. 2015. Altered machinery of protein synthesis in Alzheimer’s: from the nucleolus to the ribosome. Brain Pathol 26: 593–605.

Herzig S, Shaw RJ. 2018. AMPK: guardian of metabolism and mitochondrial homeostasis. Nat Rev Mol Cell Biol 19: 121–135.

Holtzman DM, Bales KR, Wu S, Bhat P, Parsadanian M, Fagan AM, Chang LK, Sun Y, Paul SM. 1999. Expression of human apolipoprotein E reduces amyloid-β deposition in a mouse model of Alzheimer’s disease. J Clin Invest 103: R15–R21.

Hsu C-C, Wahlqvist ML, Lee M-S, Tsai H-N. 2011. Incidence of dementia is increased in type 2 diabetes and reduced by the use of sulfonylureas and metformin. J Alzheimer’s Dis 24: 485–493.

Huang Y, Happonen KE, Burrola PG, O’Connor C, Hah N, Huang L, Nimmerjahn A, Lemke G. 2021. Microglia use TAM receptors to detect and engulf amyloid β plaques. Nat Immun 22: 586–594.

Imayoshi I, Sakamoto M, Ohtsuka T, Takao K, Miyakawa T, Yamaguchi M, Mori K, Ikeda T, Itohara S, Kageyama R. 2008. Roles of continuous neurogenesis in the structural and functional integrity of the adult forebrain. Nat Neurosci 11: 1153–1161.

Jiang Q, Lee CYD, Mandrekar S, Wilkinson B, Cramer P, Zelcer N, Mann K, Lamb B, Wilson TM, Collins JL, et al. 2008. ApoE promotes the proteolytic degradation of Aβ. Neuron 58: 681–693.

Kaise, T., Fukui, M., Sueda, R., Piao, W., Yamada, M., Kobayashi, T., Imayoshi, I., and Kageyama, R. 2022. Functional rejuvenation of aged neural stem cells by Plagl2 and anti-Dyrk1a activity. Genes Dev 36: 23–37.

Koistinaho M, Lin S, Wu X, Esterman M, Koger D, Hanson J, Higgs R, Liu F, Malkani S, Bales KR, Paul SM. 2004. Apolipoprotein E promotes astrocyte colocalization and degradation of deposited amyloid-b peptides. Nat Med 10: 719–726.

Konishi Y, Yang L-B, He P, Lindholm K, Lu B, Li R, Shen Y. 2014. Deficiency of GDNF receptor GFRn1 in Alzheimer’s neurons results in neuronal death. J Neurosci 34: 13127–13138.

Lau S-F, Chen C, Fu W-Y, Qu JY, Cheung TH, Fu AKY, Ip NY. 2020. IL-33-PU.1 transcriptome reprogramming drives functional state transition and clearance activity of microglia in Alzheimer’s disease. Cell Rep 31: 107530.

Lau S-F, Wu, W, Wong HY, Ouyang L, Qiao Y, Xu J, Lau JH, Wong C, Jiang Y, Holtzman DM, Fu AKY, Ip NY. 2023. The VCAM1-ApoE pathway directs microglial chemotaxis and alleviates Alzheimer’s disease pathology. Nat Aging 3: 1219–1236.

Li Y, Munoz-Mayorga D, Nie Y, Kang N, Tao Y, Lagerwall J, Pernaci C, Curtin G, Coufal NG, Mertens J, Shi L, Chen X. 2024. Cell Metab. 36: 1–20.

Li Y-D, Luo Y-J, Xie L, Tart DS, Sheehy RN, Zhang L, Coleman Jr LG, Chen X, Song J. 2023. Activation of hypothalamic-enhanced adult-born neurons restores cognitive and affective function in Alzheimer’s disease. Cell Stem Cell 30: 415–432.

Lin S.E., Oyama T., Nagase T., Harigaya K., Kitagawa M. 2002. Identification of new human mastermind proteins defines a family that consists of positive regulators for notch signaling. J Biol Chem 277: 50612–50620.

Liu L, Zhang K, Sandoval H, Yamamoto S, Jaiswal M, Sanz E, Li Z, Hui J, Graham BH, Quitana A, Bellen HJ. 2015. Glial lipid droplets and ROS induced by mitochondrial defects promote neurodegeneration. Cell 160: 177–190.

Lugert S, Basak O, Knuckles P, Haussler U, Fabel K, Götz M, Haas CA, Kempermann G, Taylor V, Giachino C. 2010. Quiescent and active hippocampal neural stem cells with distinct morphologies respond selectively to physiological and pathological stimuli and aging. Cell Stem Cell 6: 445–456.

Maier M, Peng Y, Jiang L, Seabrook TJ, Carroll MC, Lemere CA. 2008. Complement C3 deficiency leads to accelerated Aβ plaque deposition and neurodegeneration and modulation of the microglia/macrophage phenotype in amyloid precursor protein transgenic mice. J Neurosci 28: 6333–6341.

Mihaylova MM, Shaw RJ. 2011. The AMPK signalling pathway coordinates cell growth, autophagy and metabolism. Nat Cell Biol 13: 1016–1023.

Mary A, Eysert F, Checler F, Chami M. 2023. Mitophagy in Alzheimer’s disease: molecular defects and therapeutic approaches. Mol Psychiatry 28: 202–216.

Moreno-Jiménez E, Flor-Garceía M, Terreros-Roncal J, Rábano A, Cafini F, Pallas-Bazarra N, Ávila J, Llorens-Marteín M. 2019. Adult hippocampal neurogenesis is abundant in neurologically healthy subjects and drops sharply in patients with Alzheimer’s disease. Nat Med 25: 554–560.

Nakashiba T, Cushman JD, Pelkey KA, Renaudineau S, Buhl DL, McHugh TJ, Rodriguez Barrera V, Chittajallu R, Iwamoto KS, McBain CJ, et al. 2012. Young dentate granule cells mediate pattern separation, whereas old granule cells facilitate pattern completion. Cell 149: 188–201.

Oakley H, Cole SL, Logan S, Maus E, Shao P, Craft J, Guillozet-Bongaarts A, Ohno M, Disterhoft J, Van Eldik L, Berry R, Vassar R. 2006. Intraneuronal beta-amyloid aggregates, neurodegeneration, and neuron loss in transgenic mice with five familial Alzheimer’s disease mutations: potential factors in amyloid plaque formation. J Neurosci 26: 10129–10140.

Rath S, Sharma R, Gupta R, Ast T, Chan C, Durham TJ, Goodman RP, Grabarek Z, Haas ME, Hung WHW, et al. 2021. MitoCarta3.0: an updated mitochondrial proteome now with sub-organelle localization and pathway annotations. Nucleic Acids Res 49: D1541–D1547.

Ripa R, Ballhysa E, Steiner JD, Laboy R, Annibal A, Hochhard N, Latza C, Dolfi L, Calabrese C, Meyer AM, et al. 2023. Refeeding-associated AMPKγ1 complex activity is a hallmark of health and longevity. Nat Aging 3: 1544–1560.

Salta E, Lazarov O, Fitzsimons CP, Tanzi R, Lucassen PJ, Choi SH. 2023. Adult hippocampal neurogenesis in Alzheimer’s disease: a road map to clinical relevance. Cell Stem Cell 30: 120–136.

Sanborn MA, Wang X, Gao S, Dai Y, Rehman J. 2025. Unveiling the cell-type-specific landscape of cellular senescence through single-cell transcriptomics using *SenePy*. Nat Commun 16: 1884.

Scopa C, Marrocco F, Latina V, Ruggeri F, Corvaglia V, L Regina, F, Ammassari-Teule M, Middei S, Amadoro G, Meli G, et al. 2020. Impaired adult neurogenesis is an early event in Alzheimer’s disease neurodegeneration, mediated by intracellular Aβ oligomers. Cell Death Diff 27: 934–948.

Shao X, Li C, Lu X, Cheng J, Fan X. 2021. CellTalkDA: a manually curated database of ligand-receptor interactions in humans and mice. Brief Bioinform 22: bbaa269.

Shen H, McElhinny AS, Cao Y, Gao P, Liu J, Bronson R, Griffin JD, Wu L. 2006. The Notch coactivator, MAML1, functions as a novel coactivator for MEF2C-mediated transcription and is required for normal myogenesis. Genes Dev 20: 675–688.

Sierra A, Encinas JM, Deudero JJP, Chancey JH, Enikolopov G, Overstreet-Wadiche LS, Tsirka SE, Maletic-Savatic M. 2010. Microglia shape adult hippocampal neurogenesis through apoptosis-coupled phagocytosis. Cell Stem Cell 7: 483–495.

Straten G, Eschweiler GW, Maetzler W, Laske C, Leyhe T. 2009. Glial cell-line derived neurotrophic factor (GDNF) concentrations in cerebrospinal fluid and serum of patients with early Alzheimer’s disease and normal controls. J Alzheimers Dis 18: 331–337.

Sueda R, Imayoshi I, Harima Y, Kageyama R. 2019. High Hes1 expression and resultant Ascl1 suppression regulate quiescent vs. active neural stem cells in the adult mouse brain. Genes Dev 33: 511–523.

Tobin MK, Musaraca K, Disouky A, Shetti A, Bheri A, Honer WG, Kim N, Dawe RJ, Bennett DA, Arfanakis K, Lazarov O. 2019. Human hippocampal neurogenesis persists in aged adults and Alzheimer’s disease patients. Cell Stem Cell 24: 974–982.

van Olst L, Simonton B, Edwards AJ, Forsyth AV, Boles J, Jamshidi P, Watson T, Shepard N, Krainc T, Argue BMR, et al. 2025. Microglial mechanisms drive amyloid-β clearance in immunized patients with Alzheimer’s disease. Nat Med 31: 1604–1616.

Vázquez-Manrique RP, Farina F, Cambon K, Sequedo MD, Parker AJ, Millán JM, Weiss A, Déglon N, Neri C. 2016. Hum Mol Genet 25: 1043–1058.

Võikar V, Rossi J, Rauvala H, Airaksinen MS. 2004. Impaired behavioural flexibility and memory in mice lacking GDNF family receptor α2. Eur J Neurosci 20: 308–312.

Walgrave H, Balusu S, Snoeck S, Eynden EV, Craessaerts K, Thrupp N, Wolfs L, Horré K, Fourne Y, Ronisz A, et al. 2021. Restoring miR-132 expression rescues adult hippocampal neurogenesis and memory deficits in Alzheimer’s disease. Cell Stem Cell 28: 1805–1821.

Wang J, Gallagher D, DeVito LM, Cancino GI, Tsui D, He L, Keller GM, Frankland PW, Kaplan DR, Miller FD. 2012. Metformin activates an atypical PKC-CBP pathway to promote neurogenesis and enhance spatial memory formation. Cell Stem Cell 11: 23–35.

Wang X, Jia J. 2023. Magnolol improves Alzheimer’s disease-like pathologies and cognitive decline by promoting autophagy through activation of the AMPK/mTOR/ULK1 pathway. Biomed Pharmacother 161: 114473.

Wu L, Sun T, Kobayashi K, Gao P, Griffin JD. 2002. Identification of a family of mastermind-like transcriptional coactivators for mammalian notch receptors. Mol Cell Biol 22: 7688–7700.

Wu X, Miller JA, Lee BTK, Wang Y, Ruedl C. 2025. Reducing microglial lipid load enhances β amyloid phagocytosis in an Alzheimer’s disease mouse model. Sci Adv 11: eadq6038.

Xing C, Liu S, Wang L, Ma H, Zhou M, Zhong H, Zhu S, Wu Q, Ning G. 2024. Metformin enhances endogenous neural stem cells proliferation, neuronal differentiation, and inhibits ferroptosis through activating AMPK pathway after spinal cord injury. J Transl Med 22: 723.

Xiong X, James BT, Boix CA, Park YP, Galani K, Victor MB, Sun N, Hou L, Ho L-L, Mantero J, et al. 2023. Epigenomic dissection of Alzheimer’s disease pinpoints causal variants and reveals epigenome erosion. Cell 186: 4422–4437.

Zhang R, Boareto M, Engler A, Louvi A, Giachino C, Iber D, Taylor V. 2019. Id4 downstream of Notch2 maintains neural stem cell quiescence in the adult hippocampus. Cell Rep 28: 1485–1498.

Zhou Y, Su Y, Li S, Kennedy BC, Zhang DY, Bond AM, Sun Y, Jacob F, Lu L, Hu p, et al. 2022. Molecular landscapes of human hippocampal immature neurons across lifespan. Nature 607: 527–533.

Zimmermann HR, Yang W, Kasica N, Zhou X, Wang X, Becklman BC, Lee J, Furdui CM, Keene CD, Ma T. 2020. Brain-specific repression of AMPA α1 alleviates pathophysiology in Alzheimer’s model mice. J Clin Invest 130: 3511–3527.

