## Supplemental Data for "Activation of neurogenesis improves amyloid-β pathology and cognitive function through AMP kinase signaling in Alzheimer’s disease model mice"

**Supplemental Material**

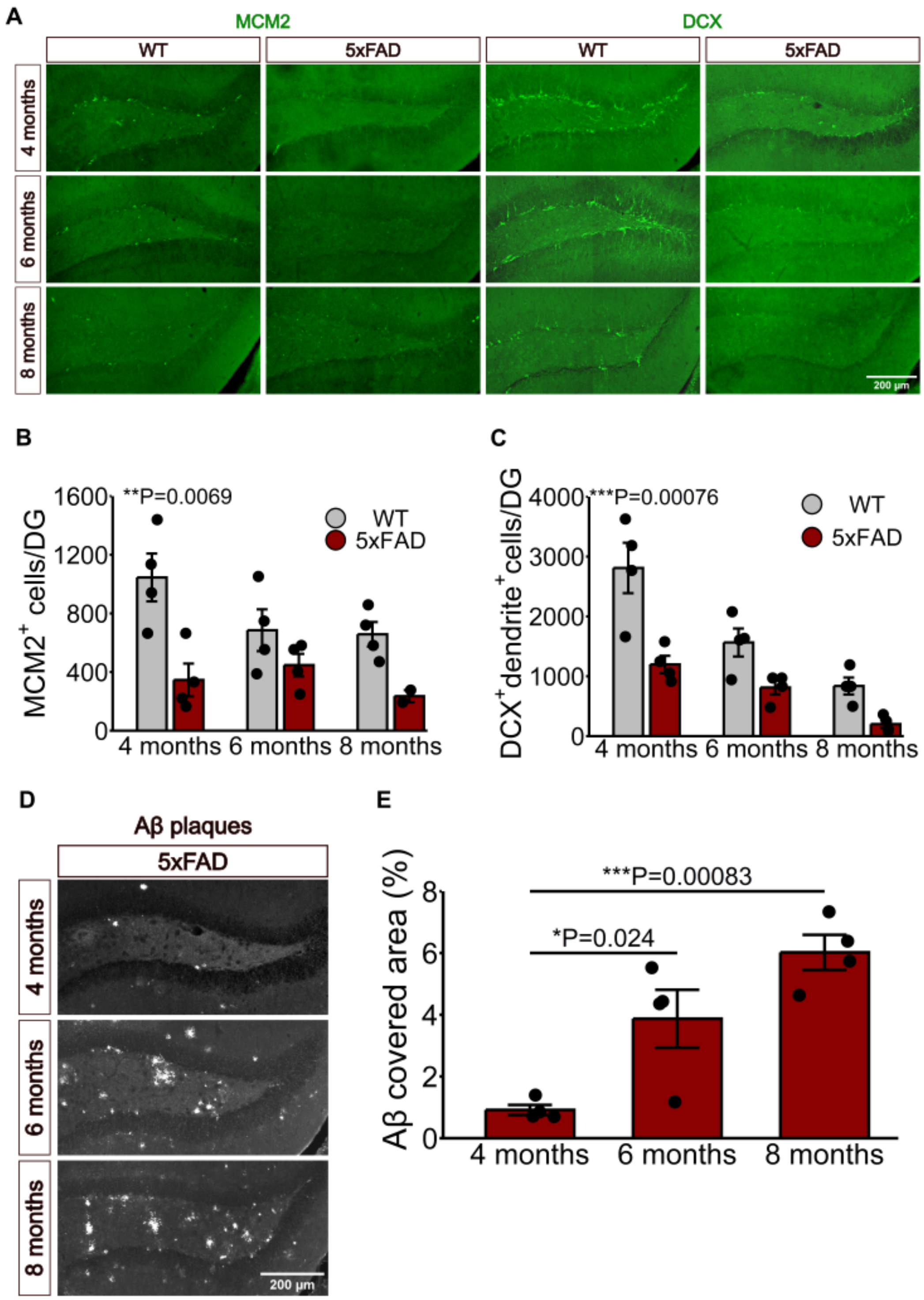


**Figure S1. Decreased neurogenesis and increased Aβ accumulation in 5xFAD mice with aging.** (A-C) Representative images (A) and quantification of $MCM2^{+}$ cells (B) and $DCX^{+}$ immature neurons (C) at 4-month, 6-month, and 8-month-old 5xFAD mice (n = 4

for each group). Scale bar: 200 μm. Values are presented as the mean ± SEM. Two-way ANOVA with Tukey's post hoc test for multiple comparisons was used in (B) and (C). (D,E) Representative images (D) and quantification (E) of the covered area of Aβ plaques at 4-month, 6-month, and 8-month-old 5xFAD mice (n = 4 for each group). Scale bar: 200 μm. Values are presented as the mean ± SEM. One-way ANOVA with Tukey's post hoc test for multiple comparisons was used in (E).

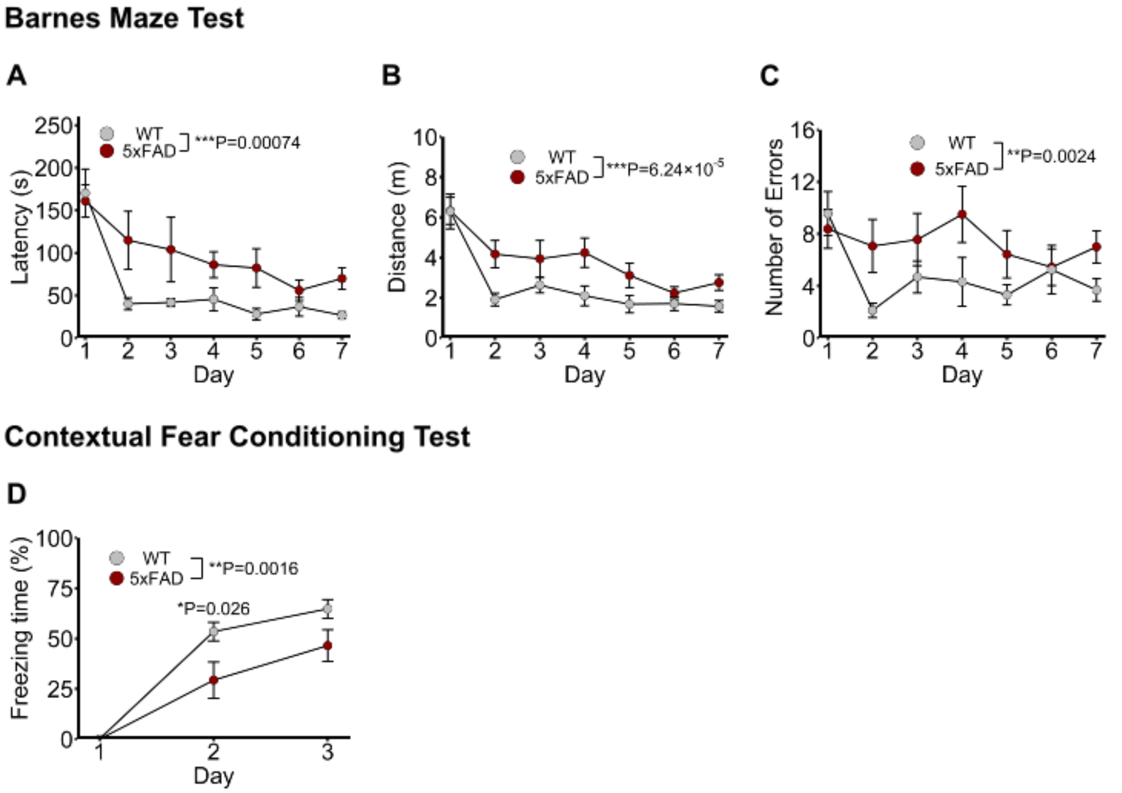


**Figure S2. 5xFAD mice show memory deficits in behavior tests.** Using wild-type and 5xFAD mice, (A) latency time, (B) distance to reach the goal hole, and (C) the number of incorrect holes visited were evaluated during the training session in Barnes maze test. (D) The percentage of time spent freezing in context A was measured during days 1-3 by the fear conditioning test. Values are presented as the mean ± SEM. *P < 0.05, **P < 0.01, ***P < 0.001. Two-way ANOVA with Tukey's post hoc test for multiple comparisons was used for statistical analysis.

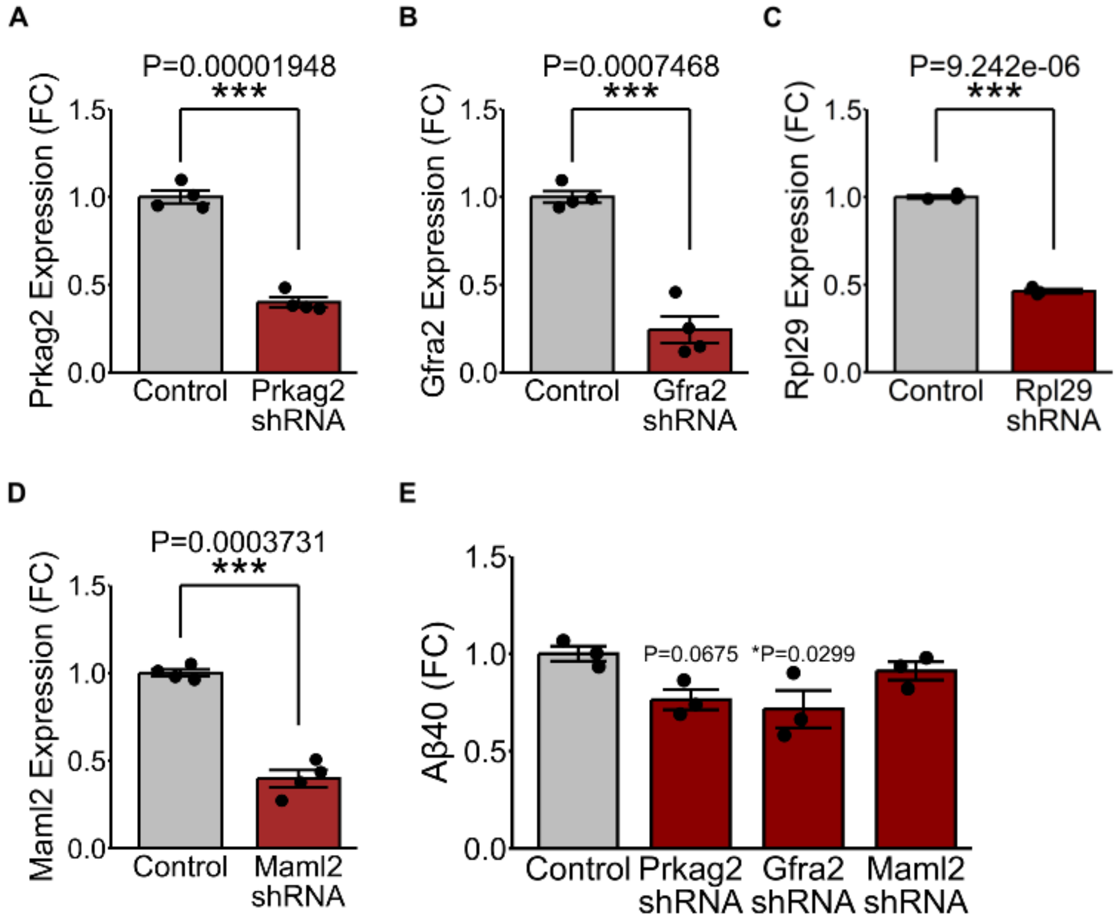


**Figure S3. Knockdown effects of *Prkag2*, *Gfra2*, *Rpl29*, and *Maml2*.** (A-D) NSCs were infected with knockdown lentivirus for *Prkag2*, *Gfra2*, *Rpl29*, or *Maml2* and subjected to qPCR analysis. (E) NSCs were infected with lentivirus for knockdown of *Prkag2*, *Gfra2*, or *Maml2*, and the culture supernatant was subjected to ELISA analysis for Aβ40. Data are presented as the mean ± SEM. *P < 0.05, **P < 0.01, ***P < 0.001. Statistical significance was assessed using Welch's *t* test for pairwise comparisons, and two-way ANOVA followed by Tukey's post hoc test for multiple group comparisons.

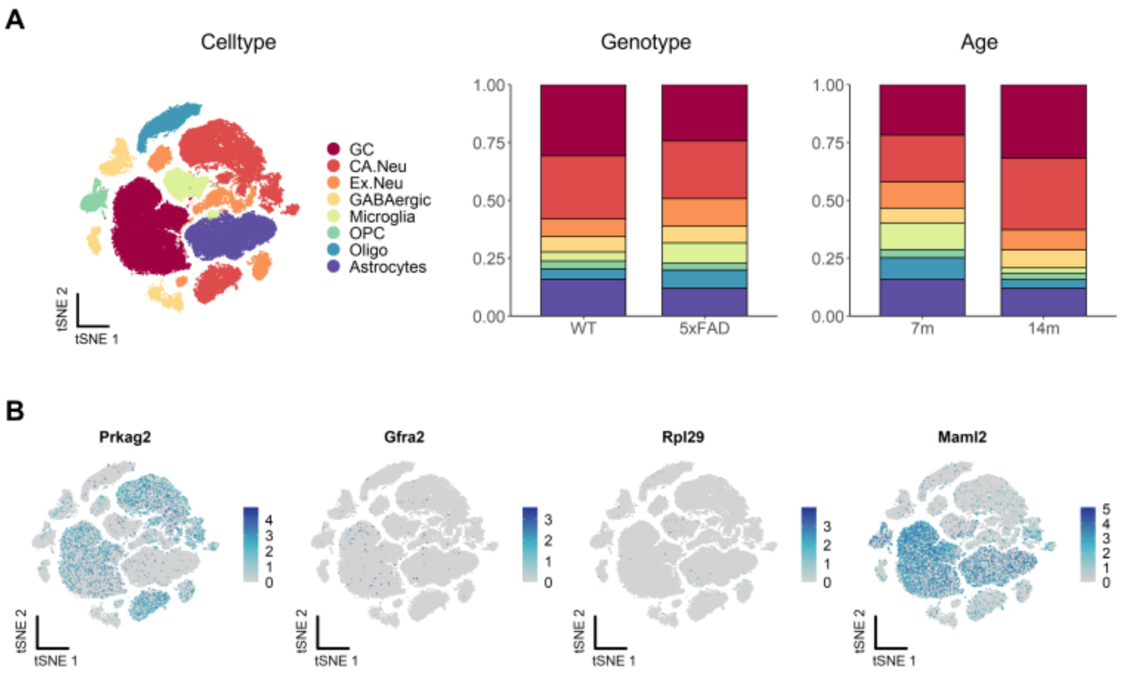


**Figure S4. Single-nucleus RNA-sequencing analysis of the hippocampus of 7- and 14-month-old wild-type and 5xFAD mice.** (A) Integrated t-SNE visualization of 7-month-old and 14-month-old wild-type and 5xFAD nuclei (left). Cell type proportions comparing wild-type (WT) and 5xFAD mice (center), and 7-month-old and 14-month-old mice (right). Different colors represent distinct cell types. GC, granule cell; CA.Neu, CA neuron; Ex.Neu, excitatory neuron; GABA, GABAergic neuron; OPC, oligodendrocyte precursor cell; Oligo, oligodendrocyte. Proportions of cellular compositions are indicated on the right. (B) Feature plots highlighting the expression levels of *Prkag2*, *Gfra2, Rpl29*, and *Maml2*.

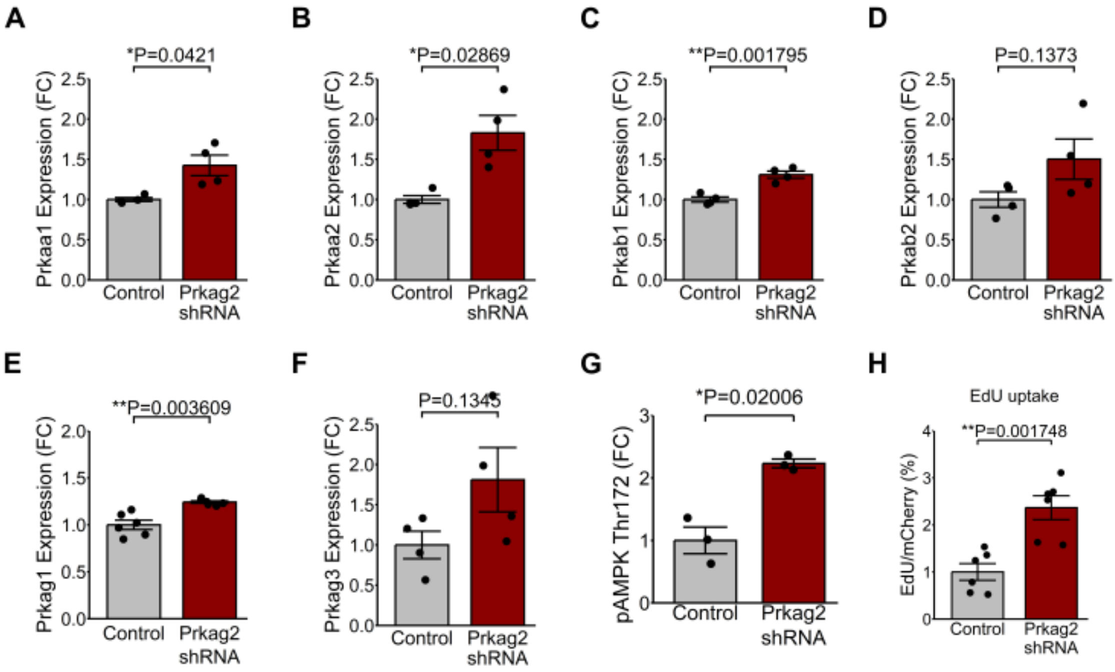


**Figure S5. The effect of *Prkag2* knockdown in NSCs.** Quiescent NSC cultures were infected with *Prkag2* knockdown lentivirus, and the effects were examined by qPCR (A-F), western blot (G), and EdU uptake (H). *Prkag1*, n = 6; Other subunit genes, n = 4. Data are presented as the mean ± SEM. *P < 0.05, **P < 0.01, ***P < 0.001. Statistical significance was assessed using Welch's *t* test for pairwise comparisons.

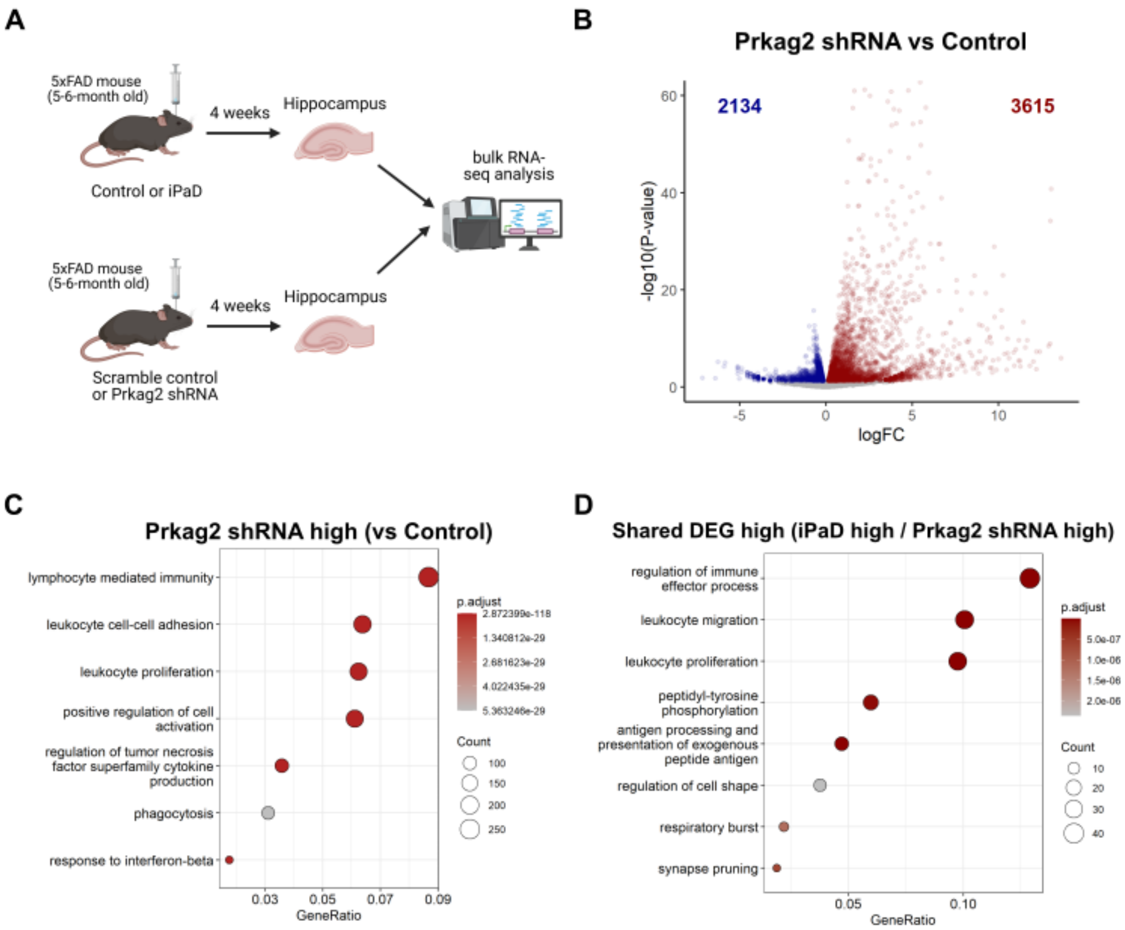


**Figure S6. RNA-seq analyses of the hippocampal dentate gyrus with *Prkag2* knockdown.** (A) Differentially expressed genes of the hippocampal dentate gyrus of 5xFAD mice between scramble control and *Prkag2* knockdown were identified and compared with those between control and iPaD treatment. Schematic was created using BioRender. (B) Volcano plot displaying differential gene expressions in the hippocampal dentate gyrus between control and *Prkag2* knockdown lentivirus-injected 5xFAD mice. (C) GO enrichment dot plot of genes upregulated in *Prkag2* knockdown lentivirus-injected 5xFAD mice compared to the control. (D) GO enrichment dot plot of genes commonly upregulated in both the iPaD and Prkag2 shRNA groups.

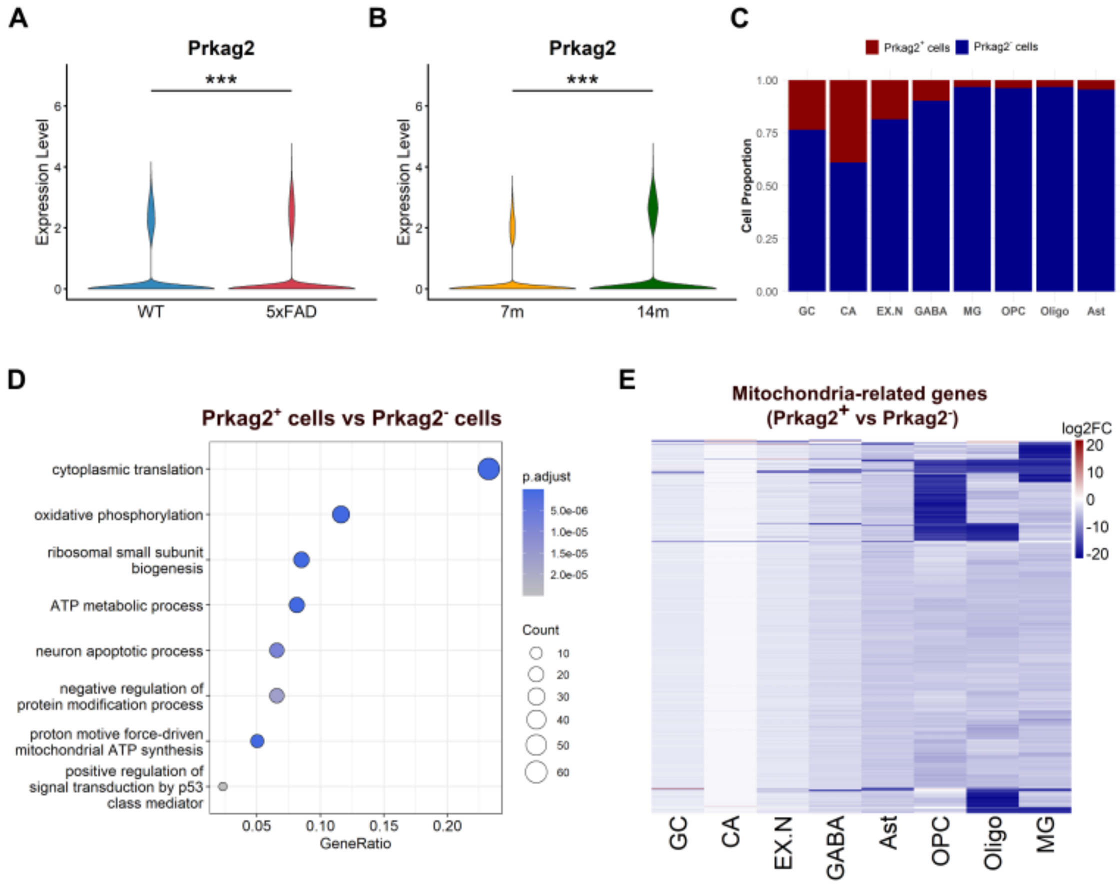


**Figure S7. Differential gene expression between *Prkag2*⁺ and *Prkag2*⁻ cells.** (A,B) Violin plots comparing *Prkag2* expression levels between wild-type (WT) and 5xFAD mice and between 7- and 14-month-old mice, based on single-nucleus RNA-seq of the hippocampus (Habib et al. 2020). (C) Stacked bar plots showing the proportions of *Prkag2*$^+$ and *Prkag2*$^-$ cells across different cell types. (D) GO enrichment dot plot of genes downregulated in *Prkag2*$^+$ cells compared to *Prkag2*$^-$ cells. (E) Heatmaps showing differential expression of mitochondria-related genes between *Prkag2*$^+$ and *Prkag2*$^-$ cells. GC, granule cell; CA, CA neuron; Ex.N, excitatory neuron; GABA, GABAergic neuron; MG, microglia; OPC, oligodendrocyte precursor cell; Oligo, oligodendrocyte; Ast, astrocyte. *P < 0.05, **P < 0.01, ***P < 0.001. Statistical significance was assessed using the two-sided Wilcoxon rank-sum test with Bonferroni correction for multiple comparisons.
